# Comparative analysis of macrophage feeder systems reveals distinct behaviors and key transcriptional shifts in chronic lymphocytic leukemia cells via coculture

**DOI:** 10.1101/2025.02.13.638101

**Authors:** Viktoria Kohlhas, Hendrik Jestrabek, Rocio Rebollido-Rios, Thanh Tung Truong, Anton von Lom, Rebekka Zölzer, Luca D. Schreurs, Duc Pham, Alexander F. vom Stein, Michael Hallek, Phuong-Hien Nguyen

## Abstract

In this brief report, we evaluated various macrophage coculture systems for their CLL-feeding potential, phagocytosis capacity, induction of treatment resistance in CLL cells, and their impact on the transcriptional profiles of CLL cells.

## Introduction

The development and progression of chronic lymphocytic leukemia (CLL) are driven not only by intrinsic properties of the leukemia cells but also by their complex interactions with the tumor microenvironment.^1^ The myeloid compartment, particularly macrophages, plays a crucial role in driving CLL progression and therapy resistance.^2,3^ Blood monocytes differentiate *in vitro* under the influence of CLL cells into nurse-like cells (NLCs), which protect leukemic cells from spontaneous apoptosis^4^ and promote multidrug resistance.^5^ Macrophage depletion *in vivo* using CSF1R blockade or liposomal clodronate, significantly reduced leukemic burden, demonstrating their important role in CLL pathogenesis.^6,7^

However, our understanding of the precise mechanisms by which macrophages promote CLL survival remains incomplete. To dissect the molecular dialog between CLL cells and macrophages, both *in vivo* models and controllable *in vitro* systems are essential. Although some macrophage-CLL coculture systems exist,^4,8–10^ systematic, simultaneous analyses of these systems are lacking. Therefore, we evaluated various macrophage coculture systems for their CLL-feeding potential, phagocytosis capacity, induction of treatment resistance, and their impact on CLL transcriptional profiles (Figure 1A).

**Figure 1.**
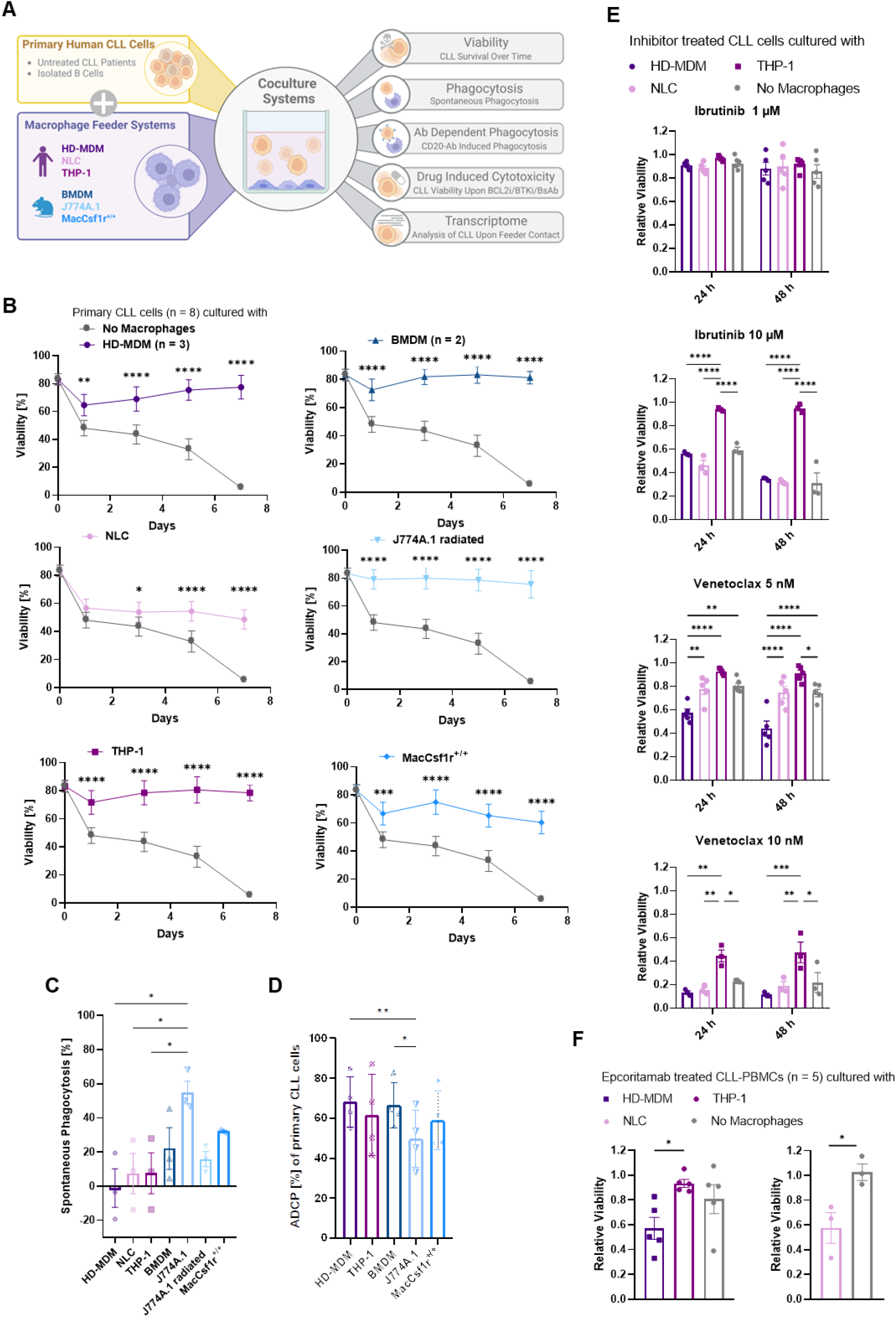
(**A**) Schematic overview of the macrophage systems and experimental readouts utilized in this study. (**B**) The viability of chronic lymphocytic leukemia (CLL) cells (n = 8) was assessed at 1, 3, 5, and 7 days in coculture with various macrophage feeder systems compared to CLL monoculture. Results are expressed as the percentage of viable CLL cells over time. All systems showed significantly prolonged CLL cell survival, analyzed with a two-way ANOVA and Šídák’s multiple comparisons test (p-values are listed in Table S12). (**C**) Phagocytosis of CLL cells (n = 3) after 18 h of coculture was quantified. Statistical analysis was performed using an ordinary one-way ANOVA followed by Tukey’s multiple comparisons test to assess differences between groups. (p-values are listed in Table S13). (**D**) Obinutuzumab-dependent phagocytosis of primary CLL cells (n = 4) was evaluated after 18 h of coculture. Statistical significance was determined via repeated measures ANOVA to account for differences between patients, followed by Tukey’s multiple comparisons (p-values are listed in Table S14). (**E**) The effect of venetoclax (5 nM, 10 nM) and ibrutinib (1 µM, 10 µM) on CLL viability was evaluated after 24 and 48 hours in both coculture and monoculture conditions. The percentage of viable cells was determined to assess treatment efficacy. Statistical testing was performed using a two-way repeated measures ANOVA test, followed by Šídák’s multiple comparisons test to account for different timepoints (p-values are listed in Table S15). (**F**) CLL cell viability was measured after 5 days of coculture of CLL PBMCs (n = 5) with different feeding systems in the presence of Epcoritamab (500 ng/ml). Viability is normalized to the viability of coculture without Epcoritamab. Statistical testing for THP-1 vs HD-MDMs vs Monoculture was done using a one-way ANOVA test followed by Holm-Šídák’s multiple comparisons test. NLC vs Monoculture was testing using a paired t-test (p-values are listed in Table S16).

## Results and Discussion

Several human and mice macrophage systems were used. Human systems included the THP-1 macrophages^10^ differentiated with phorbol-12-myristate-13-acetate, healthy donor monocyte-derived macrophages (HD-MDM) differentiated from peripheral blood mononuclear cells (PBMCs), and NLCs generated from CLL PBMCs. NLC purity was confirmed by flow-cytometry and microscopy (Figure S1). Murine systems included primary bone marrow derived macrophages^10^ (BMDMs), J774A.1^8^ and MacCsf1r^+/+^ macrophage^11^ cell lines. Whereas primary and THP-1 macrophages do not proliferate after differentiation, J774A.1 cells exhibit robust proliferation and phagocytosis. Thus, J774A.1 macrophages were γ-irradiated to halt proliferation.

All macrophage systems were cultured simultaneously with eight treatment-naïve CLL samples (Table S1). CLL viability was measured on days 0, 1, 3, 5, and 7 by flow cytometry. All macrophage systems significantly supported CLL viability throughout the 7-day period (Figure 1B) despite inter-patient variability (Figure S2A). Due to the lower NLC count, CLL viability was lowest in NLC, but this difference narrowed considerably (Figure S2B) when all macrophage systems were seeded at the same density as the average NLC count (Table S3). Moreover, fresh and thawed CLL cells showed no significant difference in viability in cocultures (Figure S3A), and the coculture systems did not induce CLL proliferation (Figure S3B). Interestingly, autologous versus allogenic CLL-NLC pairs showed no difference in viability support (Figure S4), and M2 polarization of macrophages did not enhance CLL viability beyond M0 levels (Figure S5).

To assess spontaneous phagocytic capacity, all systems were cocultured with CLL cells for 18 h, and remaining CLL cells were counted. While HD-MDMs, NLCs, and THP-1 macrophages showed modest phagocytosis, murine macrophages exhibited high rates, with non-irradiated J774A.1 cells demonstrating the highest rate, followed by MacCsf1r^+/+^ and BMDM (Figure 1C), implying that CLL cell clearance by murine macrophages should be particularly considered.

To assess antibody-dependent cellular phagocytosis (ADCP), another key function of macrophages, CLL-macrophage cocultures were exposed for 18 h to the monoclonal anti-CD20 antibody obinutuzumab. We observed no significant difference in ADCP between the systems, except for irradiated J774.A1 (Figure 1D). When combined with the higher spontaneous phagocytosis of murine macrophages, these results suggest that human macrophages are more effective in ADCP than murine ones. Altogether, these assays highlighted fundamental differences between human and murine macrophages and the importance of species-specific immune cell interactions.

Emerging evidence suggests that macrophages significantly influence treatment outcome and therapy response of CLL patients.^3^ To evaluate the suitability of the coculture systems in drug testing experiments, we treated the human macrophage-CLL cocultures with the BCL2 inhibitor venetoclax and the BTK inhibitor ibrutinib. High-dose ibrutinib (10 µM) significantly reduced the CLL cell survival cocultured with HD-MDMs or NLCs, whereas THP-1 coculture appeared to confer resistance to ibrutinib-induced CLL apoptosis, showing no remarkable viability drop (Figure 1E). Both low- and high-dose venetoclax were potent in killing most CLL cells in mono- and cocultures. However, THP-1 coculture again provided partial protection, resulting in higher CLL cell viability than other systems (Figure 1E). Aside from reduced HD-MDM viability at 10 µM ibrutinib, macrophage layers were not markedly affected by drug exposure (Figures S6). Our finding demonstrates a more significant protection of THP-1 feeder for venetoclax-treated CLL cells,^9^ suggesting that larger cohorts are needed for a definitive conclusion.

Current studies have shown promising efficacy for the bispecific antibody epcoritamab for refractory patients.^12,13^ We tested epcoritamab in a mix-culture of CLL-PBMCs with the human macrophage feeders. While HD-MDMs and autologous NLCs exhibited reduced CLL cell viability, indicating an epcoritamab-dependent T cell-mediated killing, THP-1 cells abrogated this effect, possibly due to the tumor origin of this cell line (Figure 1F).

To investigate the molecular mechanisms by which macrophages affect CLL cells, sorted CLL cells (Figure S7) of three patients after five days of coculture with HD-MDMs, THP-1 macrophages, NLCs, and BMDMs were collected and subjected to bulk mRNA sequencing. CLL cells cultured with BMDMs showed minimal transcriptional changes (Figure S8A), whereas coculture with human macrophages led to significant and distinct gene expression shifts: 192 with NLCs (Figure S8B), and 660 with THP-1 (Figure S8C), and 263 genes with HD-MDMs (Figure S8D), highlighting feeder-specific effects on CLL cells.

Gene set enrichment analysis using the hallmark collection from the Molecular Signatures Database identified eight hallmarks enriched in all three human systems (Figure 2A). Notably, “inflammatory response”, “IL2 STAT5 signaling”, “IL6 JAK STAT3 signaling”, and “KRAS signaling” were consistently upregulated in CLL cells upon coculture. Focusing on the crucial signaling pathways in B cell biology, we used a curated set of gene signatures related to signaling pathways, transcription factor regulation, and key cellular processes such as proliferation and metabolism known to influence CLL pathobiology (Supplemental Gene Set). Again, this analysis revealed upregulation of “STAT3”, “IL6”, “RAS”, and “Proliferation” across all systems (Figure 2B). JAK signaling was also elevated after HD-MDM- and NLC coculture. Additionally, pathway enrichment analyses using the Gene Ontology and Reactome databases also highlight enhanced ERK/MAPK signaling in CLL cells after cocultures (Figure S9). Altogether, these analyses highlighted the IL6/JAK/STAT3 and RAS/MAPK pathways as potential key mechanism through which macrophages may drive CLL cell survival.

**Figure 2.**
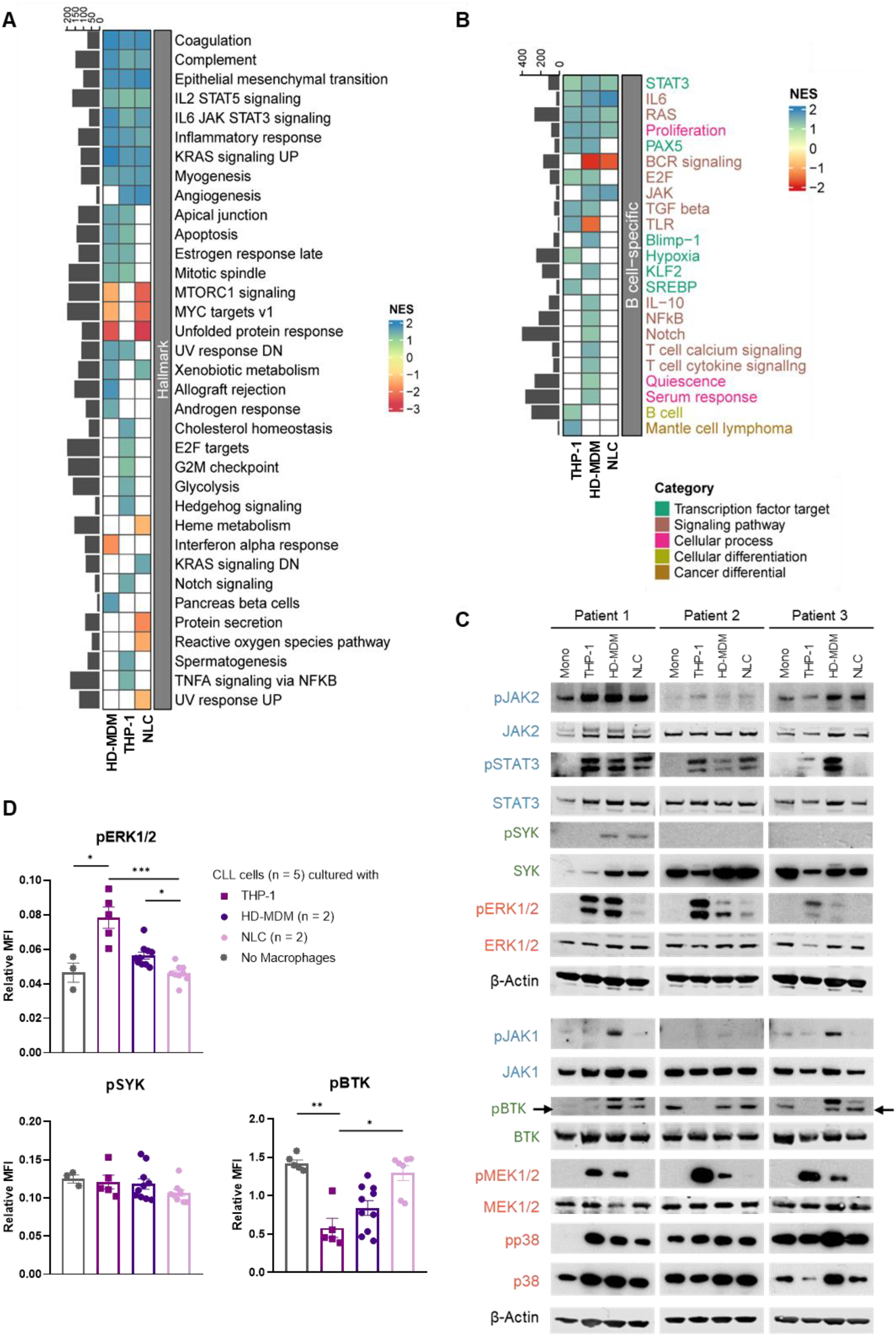
(**A**) Heatmap of enriched hallmark pathways identified by the Gene Set Enrichment Analysis (GSEA) across the different systems. The bar plots adjacent to the heatmap represent the total number of genes contributing to each enriched term, with normalized enrichment scores (NES) depicted as color gradients. NES and adjusted p-values can be found in Table S6. (**B**) Heatmap of enriched gene signatures identified through GSEA using a curated list of B-cell-specific gene sets. The gene signatures are grouped into categories, with the names color-coded according to their respective categories. The bar plots adjacent to the heatmap represent the total number of genes contributing to each enriched term, with NES depicted as color gradients. A positive score indicates enrichment in cocultured samples. NES and adjusted p-values can be found in Table S7. (**C**) CLL cells (n = 3) were lysed after 5 days of monoculture or coculture with THP-1 cells, HD-MDMs, or NLCs and analyzed by immunoblotting. Blots were probed for phosphorylated and total forms of JAK2, STAT3, SYK, ERK1/2, JAK1, BTK, MEK1/2, p38MAPK. β-Actin served as a loading control. Members of the JAK/STAT pathway are marked in blue, members of the RAS/MAPK pathway are marked in orange, members of BCR signaling are marked in green. Quantification of Western blot signals including 2 more patients is shown in Figure S10. (**D**) CLL cells (n = 5) were analyzed after 5 days in monoculture or coculture with THP-1 cells, HD-MDMs, or NLCs using phospho-flow cytometry. Levels of pSYK, pBTK, and pERK1/2 were normalized to the respective total protein levels. Statistical analysis was performed using a Kruskal–Wallis test followed by Dunn’s multiple comparisons. (p-values are reported in Table S17).

In CLL, constitutive activation of NFκB signaling in CLL cells induces IL6 production, which in turn activates the JAK/STAT pathways, creating a feed-forward loop that promotes leukemogenesis.^14^ We could confirm the activation of the JAK/STAT3 pathway by immunoblotting of independent CLL samples after cocultures (Figure 2C, Figure S10), showing increased phospho-STAT3 and phospho-JAK1 across patients in all cocultures. Phospho- and total JAK2 levels varied strongly across patients.

Elevated RAS signaling was identified in all cocultured CLL cells, revealing another pathway involved in macrophage-mediated support. Heightened RAS/MAPK activation, often driven by RAS mutations in CLL, correlates with adverse clinical features and shorter treatment-free survival.^15,16^ Immunoblotting also confirmed strongly enhanced phosphorylation of the RAS mediators MEK, ERK and p38 MAPK across patients in THP-1 and HD-MDM cocultures, despite the absence of enhanced BCR activation based on phospho-SYK and phospho-BTK levels (Figure S10), suggesting alternative upstream activation of ERK signaling such as RAS. Future studies should test whether JAK/STAT or MAPK blockade may disrupt CLL survival support.

While earlier findings have noted BCR activation upon NLC contact^17^, BCR signaling was downregulated in CLL cells cocultured with HD-MDMs and NLCs in our GSEA and in THP-1 coculture in immunoblot (Figure 2B, Figure S10). This is further confirmed by our extended analysis across multiple timepoints and culture conditions (Figure S11A), confirming enhanced BCR activation of CLL cells during NLC generation phase (Figure S11B), but unchanged or even reduced phospho-SYK and phospho-BTK during cocultures of allogeneic CLL cells with the fully-differentiated macrophages (Figure S11C). This finding illustrates the context-dependent nature of BCR regulation in CLL, and indicates that survival support provided by fully-differentiated macrophages can occur independently of BCR activation.

The CLL lymph node environment was shown to enhance “Proliferation” signature, associated with increased BCR, NFκB, and NOTCH signaling in CLL cells.^18–20^ Although we detected elevated NFκB and NOTCH signaling in CLL cells cocultured with HD-MDMs and THP-1 macrophages (Figure 2A-B), the lack of BCR activation and leukemic proliferation in our systems indicates that macrophage cocultures are insufficient to induce the prominent lymph node signatures, underscoring the collaborative role of different microenvironmental components.

## Conclusion

Collectively, all tested macrophage feeders effectively maintained CLL cell viability *in vitro*, despite notable differences in phagocytosis, ADCP capacity or drug responses. Our transcriptomic and protein-level analyses revealed that human macrophages prominently upregulate JAK/STAT and MAPK pathways in CLL cells. Given the complexity of the CLL microenvironment and the pivotal role of macrophages in disease pathogenesis, our study highlights the importance of selecting an appropriate coculture system to obtain reliable mechanistic insights. THP-1 macrophages induced the strongest transcriptional changes in the CLL cells and conferred resistance to inhibitors, warranting caution in drug-testing applications. Notably, HD-MDMs closely mirrored NLC behaviors across our assays, supporting their use as a more accessible alternative model. Cross-species systems should be interpreted with care due to elevated spontaneous phagocytosis. Finally, future studies examining macrophage polarization and the feeder system-specific differences in gene expression and cytokine production can further refine model selection and mechanistic understanding.

## Acknowledgment

We thank Daniel Bachurski, Michael Michalik, and the CLL Biobank Cologne (University of Cologne) for providing materials and patient samples, and E. Richard Stanley (Albert Einstein College of Medicine) for providing the MacCsf1r^+/+^ cell line. Figure 1A and S11A were created with BioRender.com

## Authors’ Contribution

V.K. conceptualized the study, performed experiments and analyses and wrote the manuscript. H.J. performed experiments and analyses and wrote the manuscript. R.R.R. performed computational analyses. T.T.T., R.Z., A.v.L., L.D.S., D.P., A.F.v.S. performed experiments. M.H. provided critical conceptual input and supervision. P.-H.N. conceptualized and supervised the study and wrote the manuscript. All authors read, revised and approved the manuscript.

