## Supplemental File for "Comparative analysis of macrophage feeder systems reveals distinct behaviors and key transcriptional shifts in chronic lymphocytic leukemia cells via coculture"

**Short title**: Comparative analysis of macrophage feeder systems for CLL

**Supplemental Data**

### Supplemental Methods

#### Primary CLL cells

Primary CLL cells were obtained from the patient's peripheral blood and processed by the CLL Biobank, University Hospital of Cologne. CLL cells and PMBCs were viably frozen, stored at -150 °C, and thawed upon need. Samples were only used for experimental research after written informed consent according to the Declaration of Helsinki and with Institutional Review Board approvals no. 13-091 (BioMASOTA), no. 21-1317 (SFB 1530), no. 19-1438 and no. 19-1438_1 of the University Hospital of Cologne.

#### Generation of nurse-like cells (NLCs)

In a 96-well plate, $2.5x10^6$ patient-derived PBMCs were seeded per well in a complete RPMI medium. After 14 days, cells were washed vigorously thrice with PBS to remove non-adherent cells. Purity and cell number were assessed via microscopy.

#### Isolation of murine bone marrow-derived macrophages (BMDMs)

C57B/l6 mice were kept in individually ventilated cages (IVC) in groups of a maximum of 5 individuals at the *in vivo* research facility of the CECAD Research Center, University of Cologne, under specific and opportunistic pathogen-free (SOPF) condition. All experiments were approved by the state authorities of North Rhine-Westphalia (Landesamt für Umwelt und Verbraucherschutz Nordrhein-Westfalen (LANUV)) no. 81 02.04.2019.A009. For BMDMs, femurs and tibias were harvested from 8- to 12-week-old mice. Bone ends were cut off under laminar flow. The bones were cut open on both ends to flush out the bone marrow by using a $27^{3/4}$ Ga needle and a syringe filled with sterile PBS. Afterward, marrow cells were dissociated by pressing them through a 100 µm sterile cell strainer and washed with PBS. Erythrocytes were lysed with 1 ml of ACK buffer for 3 min at RT. The cell suspension was washed twice with PBS. Cells were resuspended in DMEM (with 1% P/S, 10% FBS) containing 20 ng/ml murine M-CSF and placed in 10 cm Petri dishes to differentiate into adherent macrophages. The cells were washed, and the M-CSF-containing medium was replaced every 2 to 3 days. After a minimum of 7 days, BMDMs were harvested by trypsinization (3 ml 0.5% trypsin/EDTA for 3 min at 37 °C).

#### Generation of healthy donor monocyte-derived macrophages (HD-MDMs)

Buffy coats of healthy donors were obtained from the Transfusionsmedizin of the University Hospital Cologne, with the Institutional Review Board approvals no. 19-1559 (Buffy Coat) and no. 19-1438_1 of the University Hospital of Cologne. To isolate PBMCs from buffy coats of healthy individuals, density gradient and centrifugation with SepMate™ PBMC Isolation Tubes (STEMCELL Technologies) were performed according to the manufacturer’s instructions. In short, 15 ml of the Lymphoprep density gradient medium (STEMCELL Technologies) was carefully pipetted to the SepMate™ tube through the central hole of the insert. Subsequently,

15 ml of the buffy coat sample was diluted to a 1:1 volume ratio with PBS + 2% FBS, gently mixed, and added to the SepMate™ by pipetting it down the side of the tube. After centrifugation for 10 min at 1200 x g with the brake turned off, the top layer containing the enriched mononuclear cells was poured into a new tube. Cells were washed twice with PBS + 2% FBS and centrifuged at 300 x g for 8 min. Afterward, $1x10^8$ isolated PBMCs were seeded to a 10 cm Petri dish in differentiation medium consisting of RPMI containing 10% FBS, 1% P/S, 50 ng/ml M CSF, 1% NEAA, 1% Na Pyruvate, 2 mM. Glutamax. After 1 h of incubation, non-adherent cells were washed away with PBS, and fresh differentiation medium was added. This procedure was repeated every 2-3 days. After 7 days, cells were collected via incubation with Detachin Cell Detachment Reagent (Genlantis) for 10 min at 37 °C.

**Cocultures of CLL cells and macrophages**

Various primary and immortalized macrophage cells were chosen to compare the feeding capacity of different macrophage systems to CLL cells: BMDM, J774A.1, MacCsf1r$^{+/+}$, NLC, HD-MDM, and THP-1 macrophages. BMDMs (n = 2), NLCs (n = 6), and HD-MDMs (n = 3) were generated as described above. J774A.1 cells were Gamma radiated (40 Gy; BIOBEAM GM 8000, Gamma Service Medical) 24 h before coculture to prevent extensive proliferation and phagocytosis. THP-1 monocytes were differentiated into macrophages with 100 ng/ml PMA for 48 h. In 96-well plates, $2x10^4$ cells/well of each macrophage line were seeded, except for NLCs. NLC cell counts varied from 1483 to 4291 cells per well. To all macrophage feeder layers, $10^5$ primary CLL cells of each patient were added. To assess the viability of the CLL cells, cells were labeled with CD5-PE (clone no. REA782, Miltenyi Biotec), CD19-APC (clone no. HIB19, BioLegend), Annexin V-FITC (ImmunoTools), and DAPI (BioLegend) in Annexin V binding buffer (BD Biosciences) and subjected to flow cytometry using a MACSQuant X (Miltenyi Biotec) after 0, 1, 3, 5, and 7 days of coculture.

**Fresh CLL cell coculture**

Primary CLL cells (n = 3) were obtained from the patient's peripheral blood and processed by the CLL Biobank, University Hospital of Cologne, and immediately used for coculture. For that, $1x10^5$ CLL cells were added to $2x10^4$ THP-1 macrophages and HD-MDMs. Viability was measured after 1, 2, and 3 days via flow cytometry with Annexin V and DAPI staining, as described above. For comparison, fresh CLL cells were viably frozen and thawed again after 1 week. Cocultures were set up and measured as before.

**M2 polarization and coculture**

BMDMs were generated from wild-type mice (n = 3) as described above. After seven days of differentiation, cells were harvested using Accutase (15 min at room temperature) followed by gentle scraping. One million BMDMs per mouse were seeded into 6-well plates for either M0 control or M2 polarization. M2 differentiation was induced by treatment with 10 ng/mL IL-4 (PeproTech) and 10 ng/mL IL-13 (Miltenyi Biotec) for 48 hours. M2 polarization was assessed by flow cytometry using the following murine markers: FITC anti-mouse CD206 (BioLegend, C068C2), APC anti-mouse CD163 (BioLegend, S15049I), and PE anti-mouse CD200R (Invitrogen, OX110).

HD-MDMs were polarized to an M2 phenotype by adding 20 ng/mL human IL-4 (BioLegend) for 48 hours following M0 macrophage differentiation. Cells were detached using Accutase followed by gentle scraping. M2 polarization was analyzed by flow cytometry using the following human-specific markers: PE anti-human CD204 (Miltenyi Biotec, REA460), VioBlue anti-human CD163 (Miltenyi Biotec, REA812) and VioBlue anti-human CD206 (Miltenyi Biotec, DCN228).

THP-1 cells were differentiated into M0 macrophages with 100 ng/ml PMA for 48 h. For M2 polarization, the M0 macrophages were kept in 1 ng/ml PMA-containing medium, and 20 ng/ml IL-4 (BioLegend) and 20 ng/ml IL-13 (BioLegend) were added. M0 control macrophages were maintained in 1 ng/ml PMA-containing RPMI. The differentiation was performed for 48 h. Afterward, cells were washed twice with PBS and collected via incubation with Accutase for 10 min at 37 °C or by scraping. To label for M2 specific surface markers, cells were incubated with CD36-VioGreen (Miltenyi Biotec, REA760) and CD209-PE-Vio770 (Miltenyi Biotec, REA617) and subjected to flow cytometry (MQX).

THP-1 macrophages, BMDMs (n = 2), and HD-MDMs (n = 2) in M0 and M2 polarization were seeded for coculture with CLL cells (n = 3) in a 96-well plate and the CLL cell viability was assessed over time as described above.

**Proliferation assay**

$2x10^4$ Macrophage feeder systems (NLC (n = 2), HD-MDM (n = 2), THP-1, BMDM (n = 2), MacCsf1r$^{+/+}$, and irradiated J774A.1) were seeded in 96-well flat-bottom plates as described before. CLL patient cells (n = 5) and JVM-3 controls were washed twice with PBS, stained with 0.25 µM CFSE (1:2000 dilution; BD Horizon) for 15 min at 37 °C, quenched with cold FCS, and washed twice with RPMI. After 5 days, cells were stained with DAPI and analyzed by flow cytometry to assess proliferation via CFSE dilution. Expansion Index (EI) was calculated as the total cell count across all dye-dilution generations ($\sum N_i$) divided by the theoretical starting cell number ($\sum[N_i/2^i]$, where $N_i$ is the number of cells in generation i), yielding the fold-expansion of the population.

**Spontaneous phagocytosis**

To measure spontaneous phagocytosis of CLL cells by the macrophage feeder layers, macrophages and CLL cells (n = 3) were seeded as in coculture experiments (described above) in technical triplicates. After 18 h of coculture, CLL cells were identified via CD19-APC staining and counted via flow cytometry (MACSQuant X). This method quantifies the non-phagocytosed CLL cells remaining in the co-cultures, indirectly reflecting the extent of phagocytosis. The mean of technical triplicates was calculated and normalized to monoculture counts.

**Antibody-dependent cellular phagocytosis (ADCP)**

In 96-well plates, $1x10^4$ macrophages per well were seeded according to the cell characteristics described above. $5x10^4$ primary CLL cells (n = 4) were added to each feeder layer. Obinutuzumab (GA101, Roche, Basel, Switzerland) was added to half of the samples. All samples were prepared in quintuplets. After 18 h, plates were put at 4 °C for 30 min to stop the ADCP process. Primary CLL cells were labeled with CD19-APC (clone no. HIB19, BioLegend) before flow cytometry. CLL cells were measured with a MACSQuant VYB (Miltenyi Biotec). The ratio between spontaneous phagocytosis (no Obinutuzumab) and Obinutuzumab-treated wells was calculated.

**Inhibitor treatment**

In a 96-well plate, $2x10^4$ cells of HD-MDMs, NLCs, and THP-1 cells were seeded per well. NLCs were obtained in a 96-well plate as described above. $1x10^5$ primary CLL cells were added to all feeder systems as well as monoculture control. Either 1 µM ibrutinib and 5 nM venetoclax [CLL cells (n = 5)], 10 µM ibrutinib and 10 nM venetoclax [CLL cells (n = 3)], or DMSO was added to the wells, and viability was measured after 24 h and 48 h by flow cytometry as described above. The viability was normalized to untreated cocultured controls.

**XTT assay**

HD-MDMs, THP-1 cells, and NLCs ($2x10^4$ cells per well for HD-MDMs and THP-1; varying numbers for NLCs) were seeded in 96-well plates as previously described. Following treatment with venetoclax and ibrutinib, cell viability was assessed using the XTT Cell Viability Kit (Cell Signaling Technology) according to the manufacturer's instructions at the indicated time points.

**Bispecific antibody treatment**

$10^5$ CLL PBMCs from 5 patients were seeded on top of $2x10^4$ cells of HD-MDMs and THP-1 per well in a 96-well plate. NLCs were obtained as described above from 3 patients and $10^5$ freshly thawed, autologous CLL PBMCs were seeded on top. Cocultures were then treated with Epcoritamab (500 ng/ml) for 5 days and viability was subsequently assessed by flow cytometry. The viability was normalized to untreated cocultured controls.

**Next generation mRNA sequencing**

Cocultures of CLL patient cells (n = 3) with MDMs, THP-1 macrophages, NLCs, and BMDMs were set up as described above. Feeder macrophages were seeded to a final density of $5.5x10^5$ cells per 35x10 mm dish. $2.75x10^6$ CLL cells were seeded per dish. After 5 days of coculture, the suspension cells of cocultures were collected. CLL cells contained in the suspension were purified using CD19 MicroBeads (Miltenyi Biotec), which magnetically label $CD19^+$ cells and separate them via the magnetic field of a MACS Separator. The purification was performed using MACS LS Columns following manufacturer's instructions.

For the isolation of RNA for Next Generation mRNA sequencing (NGS), the RNeasy Plus Mini Kit (Qiagen) was used according to the manufacturer's instructions. A paired-end (2x100bp) mRNA sequencing run with poly-A selection was performed at the Cologne Center for Genomics (CCG). The Illumina TruSeq stranded NEB protocol was used according to the manufacturer's instructions for sample preparation. The sequencing was run on a NovaSeq 6000 Sequencing system (Illumina).

**Differential expression analysis**

Raw data files from all the aforementioned systems were used to map reads to the human reference genome GRCh38.p14 (GENCODE release 44) using the STAR program version 2.7.10a[1]. The reads were processed as paired-end reads, followed by quality control analysis. Subsequently, read summarization was performed using the featureCounts program version 2.0.3[2], generating count matrices as output.

To ensure the sequenced CLL cells were not confounded by inherited macrophages, we applied a filter for macrophage-specific genes. First, single cell RNA sequencing (scRNA-seq) data was retrieved from[3], focusing on the blood and immune cell transcriptomes. B-cell- and macrophage cell types were selected, resulting in 348 and 503 genes with elevated expression, respectively. Overlapping genes between the two cell types were excluded, and only genes with elevated expression categorized as "group enriched" in the macrophage cluster were retained. Next, the resulting list of 34 genes was compared with the BMDM system, and intersecting genes were removed. The remaining 24 genes were manually curated to ensure only macrophage-specific genes were included. These genes were subsequently removed from the genome before further data processing and can be found in Table S4.

Next, genes with low expression values were filtered out, retaining only those with sufficiently high counts for statistical analysis. Normalization factors were calculated to scale the raw library sizes using the trimmed mean of M-values (TMM) method. $\text{Log}_2$ counts per million (CPM) were computed to quantify gene expression levels. Differentially expressed genes were identified using the voom method with sample quality weights from the limma R package[4] based on the criteria of a q-value $\leq 0.05$ and $1 \leq \text{log2-FC} \leq -1$.

Volcano plots were generated using the ggplot2 package[5].

**Gene set enrichment analysis (GSEA)**

GSEA[6] of the pre-ranked gene list was performed using the clusterProfiler R package[7], implementing the GSEA algorithm and the fgsea R package[8] for a faster GSEA approach.

Two separate analyses were conducted: one using the Hallmark collection from the Molecular Signatures Database (MsigDB v2023.1.Hs) and another using a newly updated 2022 version of the curated list of B cell-specific signatures obtained from the Signature Database (https://lymphochip.nih.gov/signaturedb, see Supplemental Data Set). Results from both analyses were visualized using the ComplexHeatmap R package[9].

**Western Blot**

Cocultures of CLL patient cells (n = 5) with HD-MDMs, THP-1 macrophages and NLCs, were set up as described above. HD-MDM and THP-1 macrophages were seeded to a final density of $2.4 \times 10^6$ cells per 35x10 mm dish. $1.2 \times 10^7$ CLL cells were added per dish. After 5 days of coculture, the suspension cells of cocultures were collected. CLL cells contained in the suspension were purified using CD19 MicroBeads (Miltenyi Biotec) and MACS LS Columns following manufacturer's instructions.

Cell pellets were lysed in 30 µL RIPA buffer (Cell Signaling Technologies) supplemented with protease inhibitor (Complete Mini, EDTA-free; Roche) and phosphatase inhibitor (PhosSTOP; Roche), followed by incubation on ice for 60 minutes. Lysates were centrifuged at 16000 x g for 20 minutes at 4 °C, and protein concentrations were determined using the Pierce™ BCA Protein Assay Kit (Thermo Fisher Scientific), according to the manufacturer’s instructions.

For gel electrophoresis, 20 µg of protein per sample was denatured at 72 °C for 10 minutes in 1x LDS sample buffer with 1x reducing agent (Thermo Fisher Scientific). Samples and a prestained protein marker (Bio-Rad) were loaded onto 4–12% Bis-Tris gels and run using the NuPAGE™ system (Thermo Fisher Scientific) at 150 V in MOPS-SDS running buffer with 400 µL NuPAGE Antioxidant.

Proteins were transferred to nitrocellulose membranes (Amersham Biosciences) using the Trans-Blot SD Semi-Dry Transfer Cell (Bio-Rad) at 45 mA per gel for 2 hours. Membranes were blocked for 1 hour at room temperature (RT) with 5% milk blocking buffer (for HRP detection) or Intercept™ (TBS) Blocking Buffer (LI-COR, for fluorescence detection), then incubated overnight at 4 °C with primary antibodies diluted per manufacturer’s recommendations. The next day, membranes were washed 3x for 10 minutes with TBS-T.

For chemiluminescence detection, membranes were incubated for 1 hour at RT with HRP-conjugated secondary antibodies (1:10,000 in milk blocking buffer), washed, and developed using either WesternBright™ ECL substrate (Advansta) or SuperSignal™ West Femto substrate (Thermo Fisher) for low-abundance targets. Signal was detected using Super Rx X-ray film (Fujifilm).

For fluorescence detection, fluorophore-conjugated secondary antibodies (1:10,000 in Intercept buffer) were used, and membranes were imaged using the LI-COR ODYSSEY CLx system.

Band intensities were quantified using ImageJ software. Protein expression was normalized to loading controls from the same samples via densitometry.

Primary and secondary antibodies are listed in Table S8 and S9.

**BCR activity flow cytometry**

NLC samples were generated as described above. After 14 days, nonadherent lymphoid cells were removed, and the adherent NLC layer was washed twice with PBS. The nonadherent cells and PBS wash fractions were pooled and subjected to $CD19^+$ magnetic separation (as previously described) to isolate CLL cells. Isolated CLL cells were rested on ice for at least 1 hour. Half of the cells were left untreated, while the remaining half were stimulated via the B cell receptor (BCR) by incubation with 20 µg/mL Goat $F(ab')_2$ Anti-Human IgM-UNLB (Southern Biotech) for 10 minutes. Cells were washed twice with PBS to remove unbound IgM.

Cocultured CLL cells were harvested after 2, 5, and 14 days of coculture with THP-1 macrophages, HD-MDM, or NLCs.

For all samples, cells were fixed and permeabilized using the IntraPrep™ Permeabilization Reagent Kit (Beckman Coulter), according to the manufacturer's instructions, with a modified protocol using 50 µL of Reagent 2 instead of 100 µL and intracellular antibody incubation overnight at 4 °C. On the following day, cells were washed twice with PBS and analyzed by flow cytometry.

Antibodies are listed in Table S10.

**Statistical analysis**

For statistical analysis, one- and two-way ANOVA tests followed by multiple testing were used, as indicated in the figure legends. All statistical differences were calculated with the GraphPad Prism 8 software. Statistical significance was assumed at p values ≤ 0.05. Only significant changes are indicated in the figures. Asterisks were used to visualize significance:

* = p ≤ 0.05, ** = p ≤ 0.01, *** = p ≤ 0.001, **** = p ≤ 0.0001

**Data Availability**

The raw- and processed RNA sequencing data generated in this study have been deposited in the ArrayExpress database under accession code E-MTAB-15353.

### Supplemental Figures and Legends

#### Figure S1

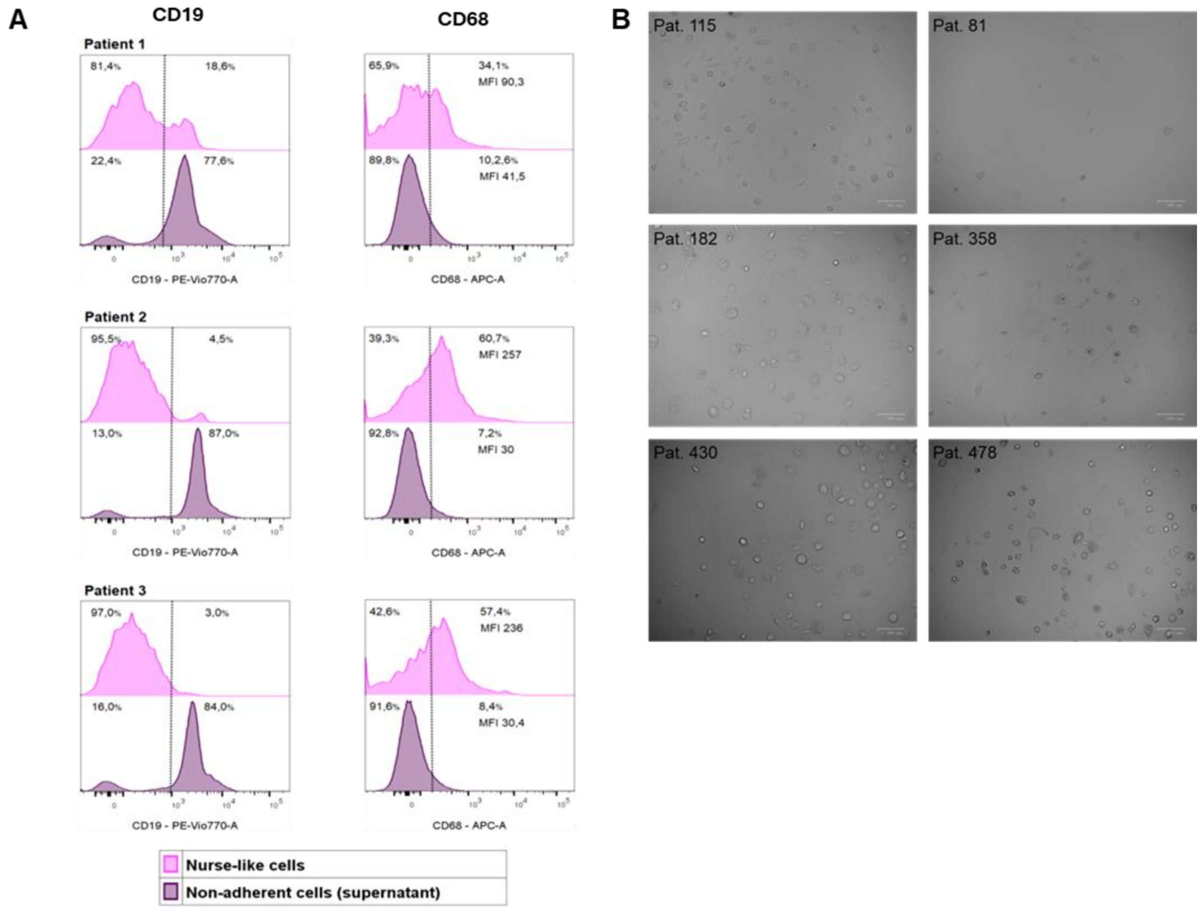

(**A**) After 14 days of CLL PBMC culture, cells were washed to remove non-adherent cells. The non-adherent cells (supernatant) and the remaining adherent NLC layers were analyzed by flow cytometry. Representative modal histograms of CD19 and CD68 expression demonstrate efficient depletion of CLL cells and enrichment of NLCs in the adherent layers following washing.

(**B**) Light microscopy images of NLC feeder layers from 6 different CLL patient cultures post-washing confirm morphological purity of the adherent NLC population. Those NLC feeder cells were used for viability experiments in Fig. 1B.

**Figure S2**

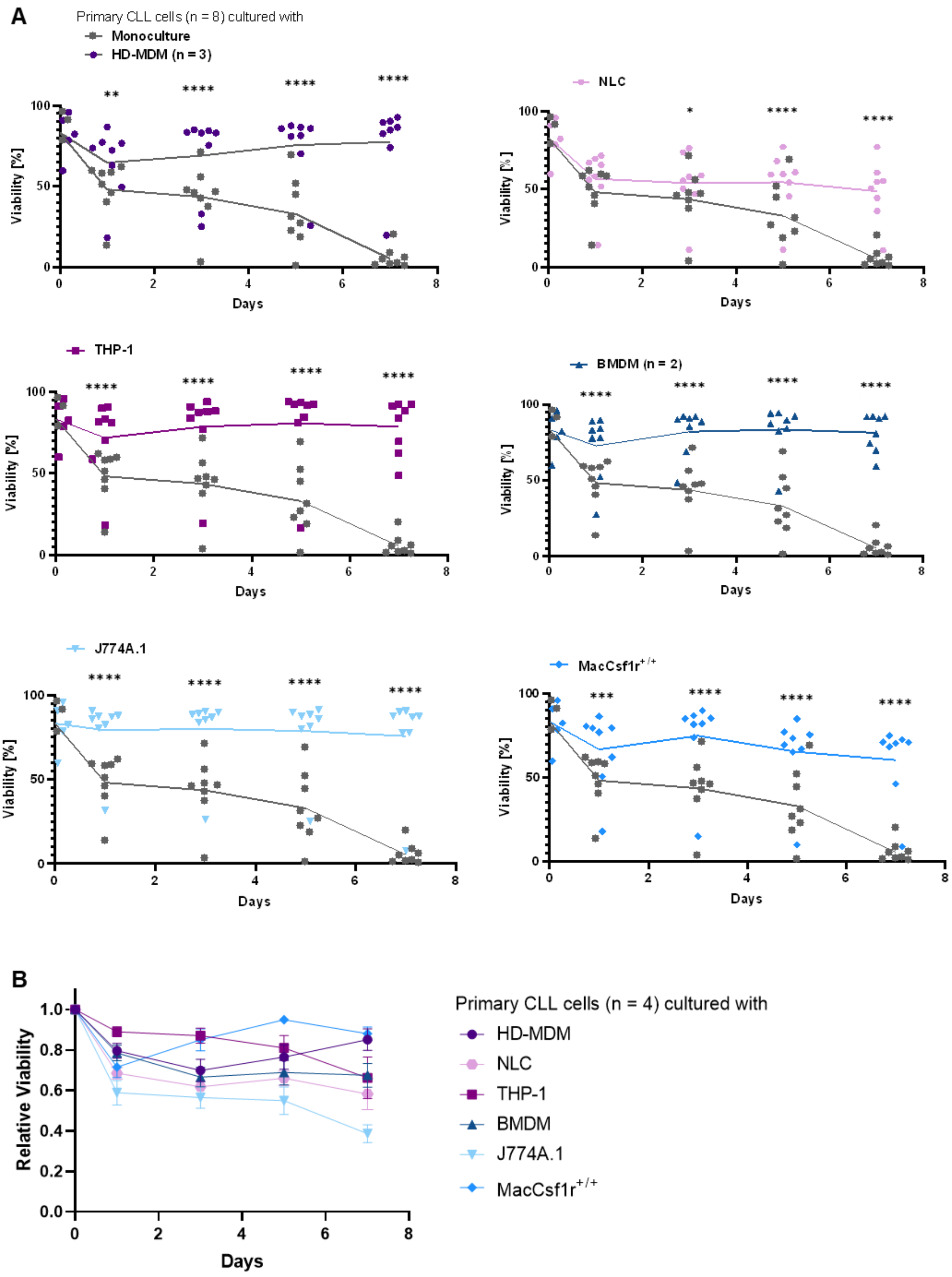

(**A**) Visualization of individual data points corresponding to the CLL viability analysis shown in Fig. 1B.

(**B**) NLCs were quantified after 14 days of CLL PBMC culture, and the average NLC cell number was calculated (2,680 cells). This number was used to normalize seeding of various macrophage types, including HD-MDMs, THP-1 cells, BMDMs, J774A.1 cells, and MacCsf1r$^{+}$/$^{+}$ cells. CLL cell viability was assessed by flow cytometry at 0, 1, 3, 5, and 7 days of coculture.

**Figure S3**

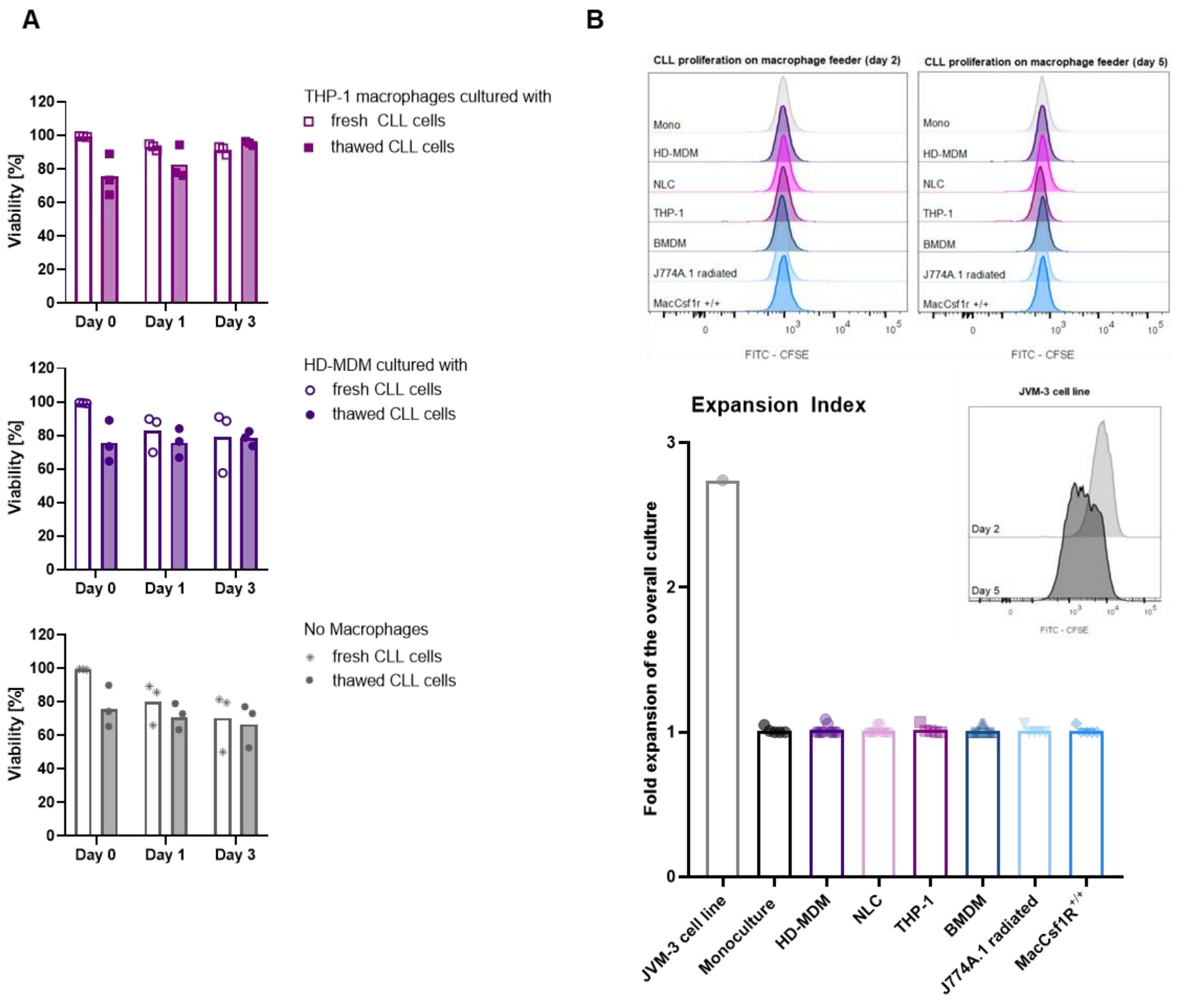

(**A**) CLL cells from 3 patients were either used freshly or frozen and thawed after one week, then cultured with THP-1 macrophages and HD-MDM. Viability was assessed over 1, 2, and 3 days.

(**B**) CFSE-labeled CLL cells (n = 5) were cultured in monoculture or in coculture with various macrophage feeder layers. Proliferation was quantified by flow cytometric analysis of CFSE dilution, and expansion indices were calculated (>1 indicating proliferation). JVM-3 cells served as positive controls.

**Figure S4**

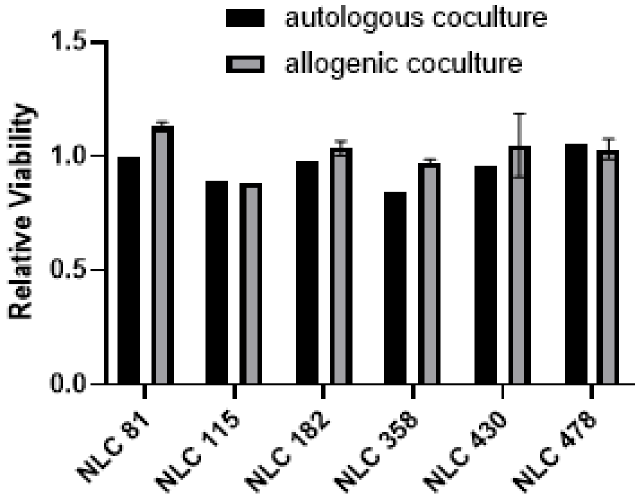

CLL cell viability normalized to the mean of all patients after 5 days of coculture with either autologous (n = 1) or allogenic (n = 2) NLCs.

### Figure S5

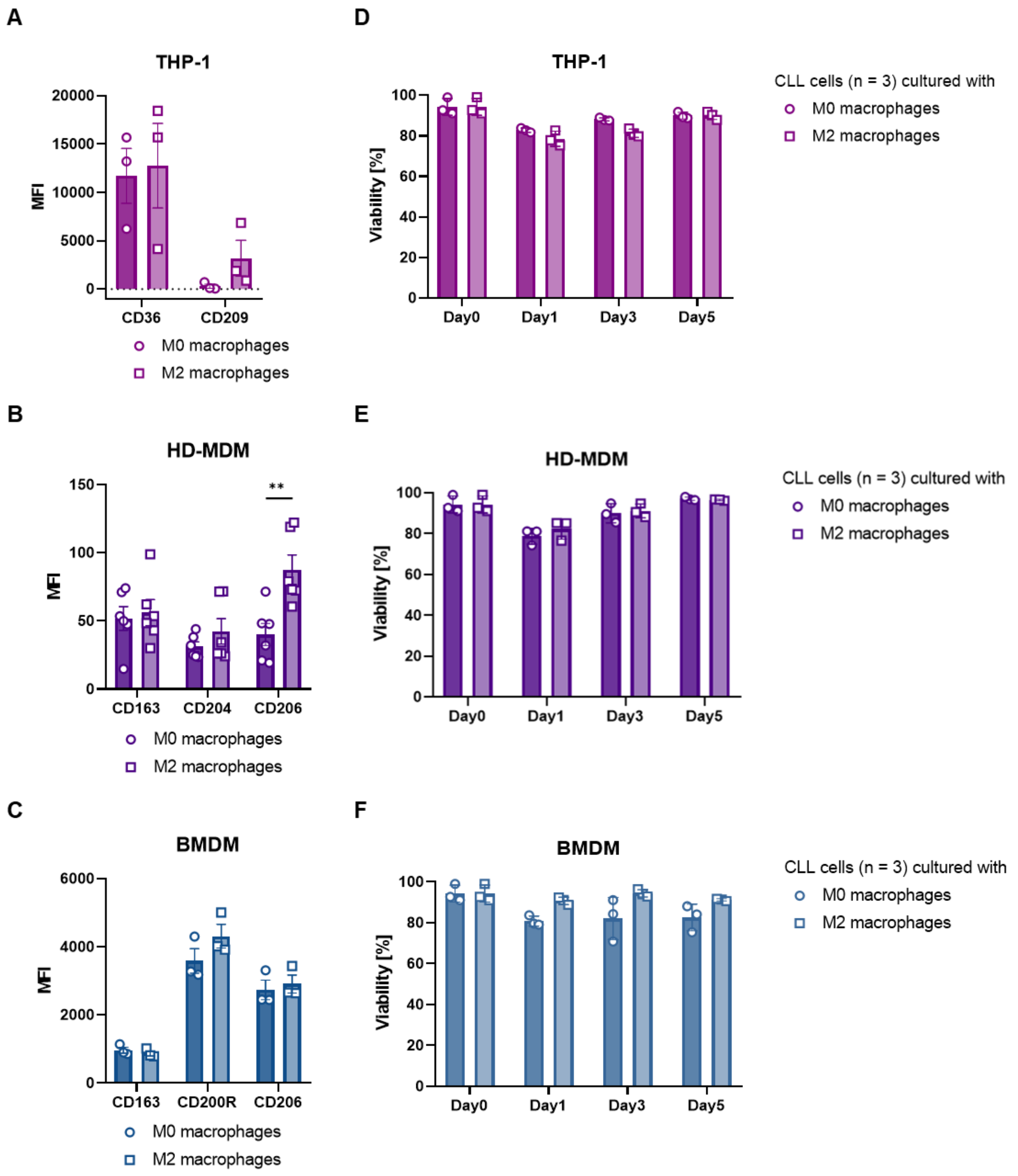

(**A-C**) Flow cytometric analysis of M2 marker expression in THP-1 macrophages (**A**), HD-MDMs (**B**), and BMDMs (**C**) after M2 polarization. Statistical analysis was performed using a two-way RM ANOVA with uncorrected Fisher's LSD multiple testing. (p-values are listed in Table S18)

(**D-F**) THP-1 cells, HD-MDMs, and BMDMs were cultured under naïve (M0) conditions or polarized toward an M2 phenotype. Subsequently, primary CLL cells (n = 3) were cocultured with different macrophage conditions, and CLL viability was assessed over 5 days by flow cytometry. No significant difference in viability support was observed between M0 and M2 macrophages, indicating comparable feeder capacity.

**Figure S6**

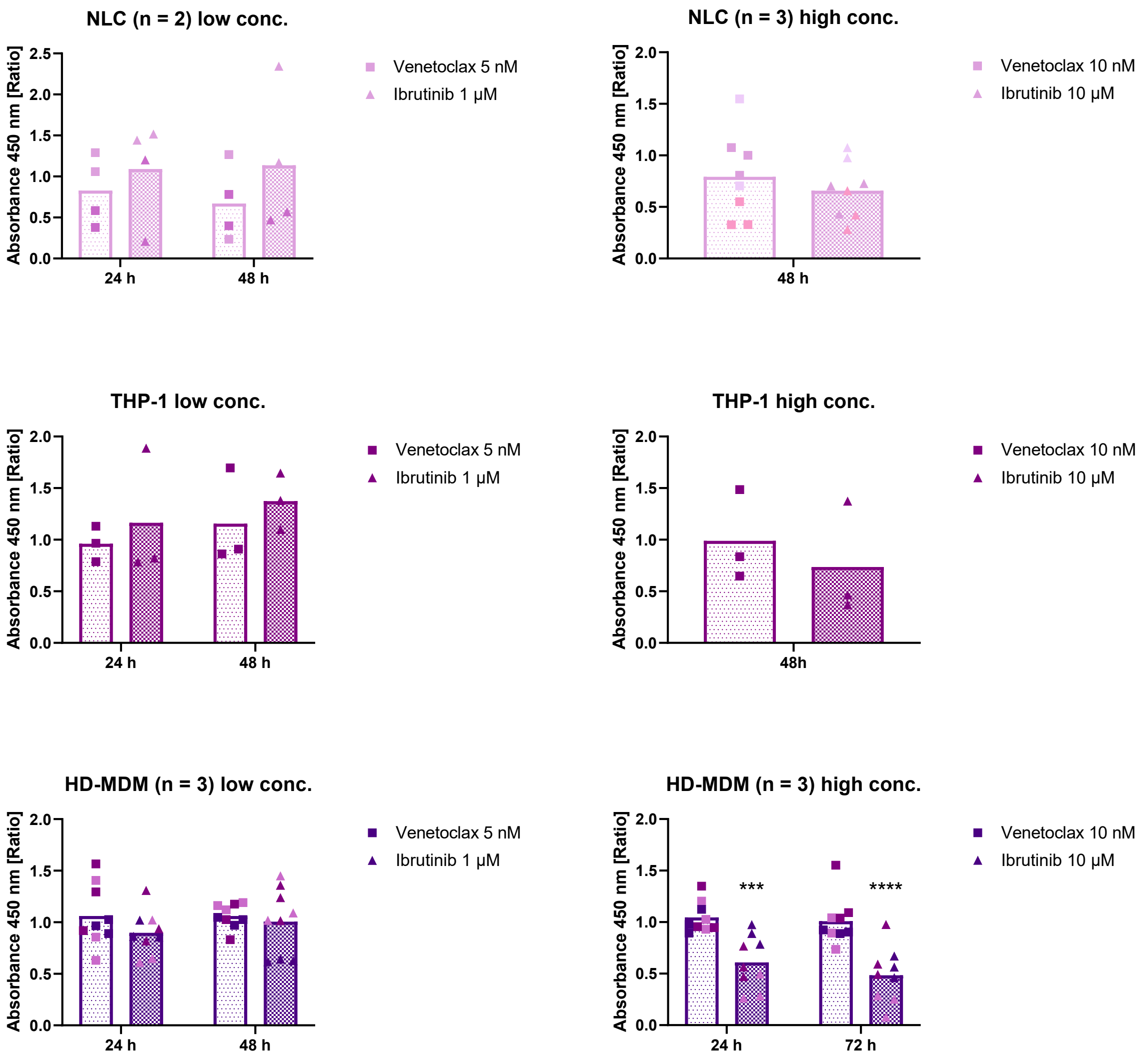

XTT assays of HD-MDMs, THP-1 macrophages, and NLCs mono-feeder layers after treatment with venetoclax ([5 nM = low conc.] [10 nM = high conc.]) and ibrutinib ([1 µM = low conc.][10 µM = high conc.]) at different time points. The measured absorbance was normalized to the untreated control. Different donors (n) are represented by differentially colored data points. Statistical analysis was performed with an ordinary one-way ANOVA (one time point) or two-way ANOVA (two time points) with a subsequent Tukey's multiple comparison. (p-values are listed in Table S19).

### Figure S7

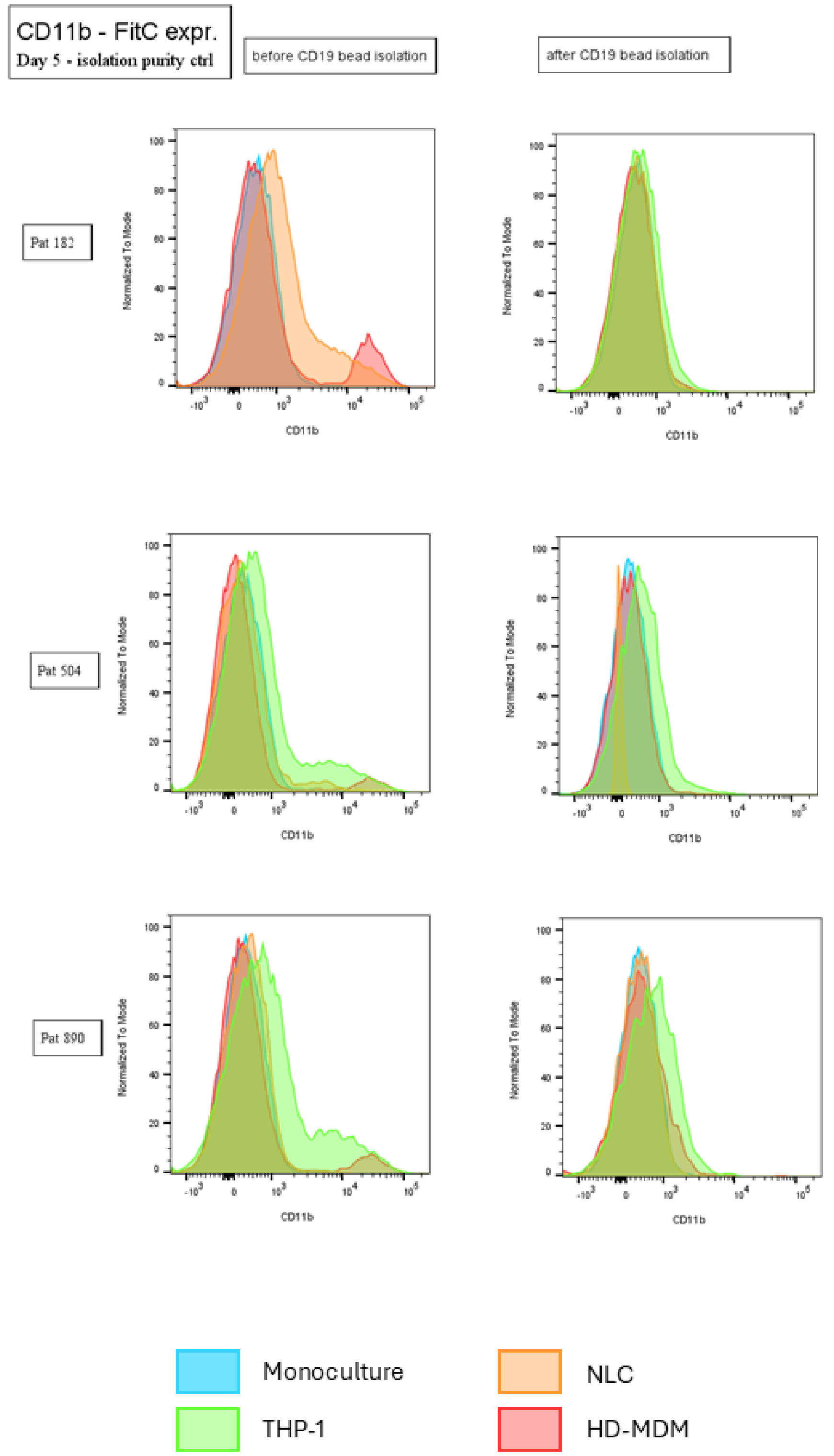

**Pat 182**

Before isolation:

| Feeder | CD11b+ (%) |
|---|---|
| NLC | 16,1 |
| Monoculture | 1,1 |
| HD-MDM | 13,9 |

After isolation:

| Feeder | CD11b+ (%) |
|---|---|
| THP-1 | 1,03 |
| NLC | 0,49 |
| HD-MDM | 0,83 |
| Monoculture | 0,24 |

**Pat 504**

Before isolation:

| Feeder | CD11b+ (%) |
|---|---|
| THP-1 | 12,4 |
| NLC | 4,54 |
| HD-MDM | 5,68 |
| Monoculture | 1,02 |

After isolation:

| Feeder | CD11b+ (%) |
|---|---|
| THP-1 | 2,38 |
| NLC | 0,043 |
| HD-MDM | 0,7 |
| Monoculture | 0,34 |

**Pat 890**

Before isolation:

| Feeder | CD11b+ (%) |
|---|---|
| THP-1 | 16,1 |
| NLC | 1,19 |
| HD-MDM | 6,97 |
| Monoculture | 0,71 |

After isolation:

| Feeder | CD11b+ (%) |
|---|---|
| THP-1 | 2,58 |
| NLC | 0,33 |
| HD-MDM | 1,34 |
| Monoculture | 0,22 |

CD11b expression was analyzed by flow cytometry in cells collected from the supernatant when washing CLL cells off the feeder layer on day 5 of coculture. Histogram plots show a representative example from three independent samples, demonstrating $CD11b^+$ myeloid cells before and after $CD19^+$ cell depletion using magnetic bead isolation. The data confirm effective removal of contaminating myeloid cells that may have detached from the feeder layer.

**Figure S8**

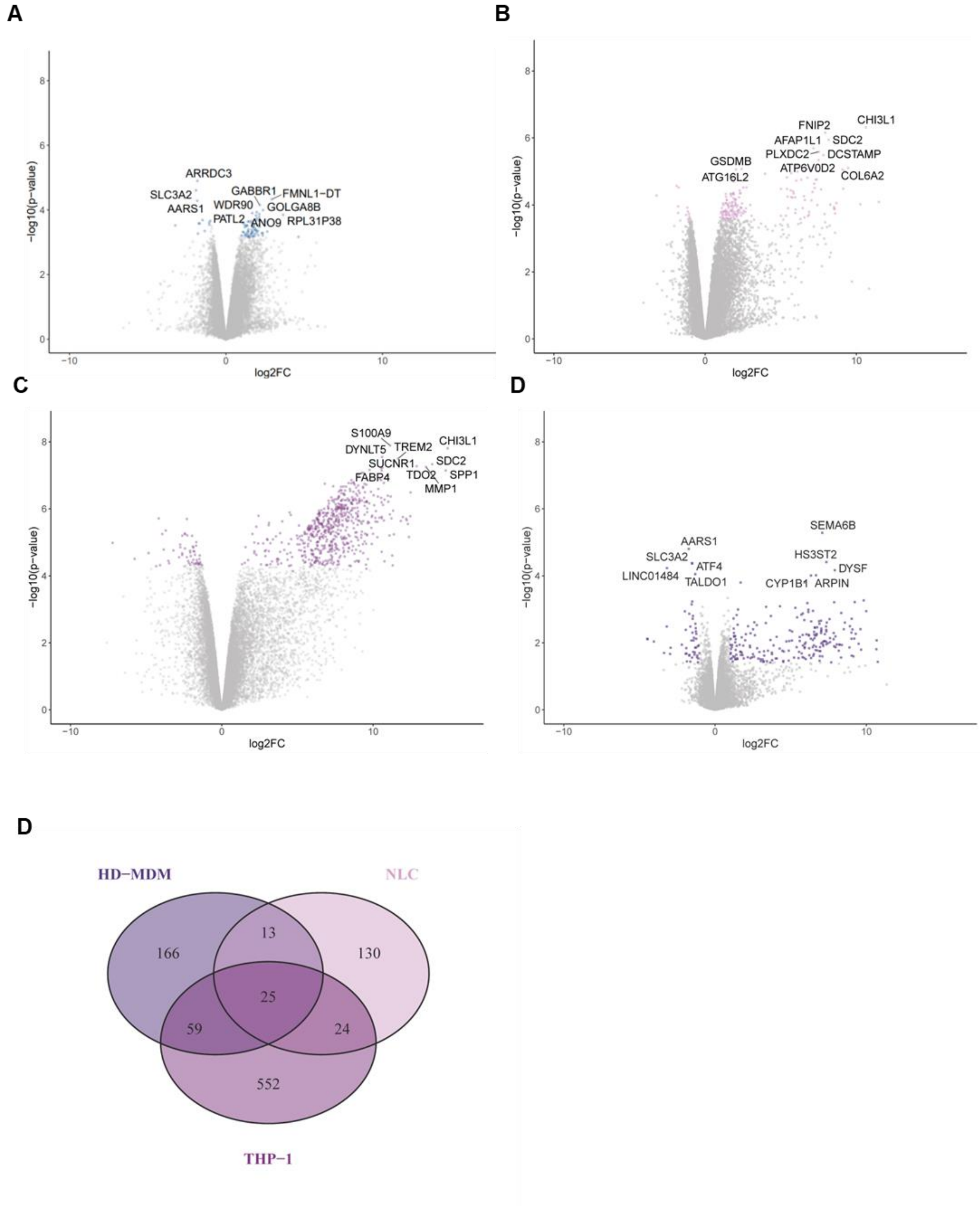

Volcano plots depicting the log2 fold-change (FC) in gene expression from bulk RNA-seq analysis of CLL cells cocultured for 5 days with (**A**) BMDMs, (**B**) NLCs, (**C**) THP-1 macrophages and (**D**) HD-MDMs compared to CLL cells in monoculture. The y-axes represent the –log10 p-value. Differentially expressed genes (DEGs) are highlighted in color (q-value < 0.05 and 1 ≤ log2-FC ≤ −1). The top 10 genes ranked by q-value in each coculture system are labeled.

**Figure S9**

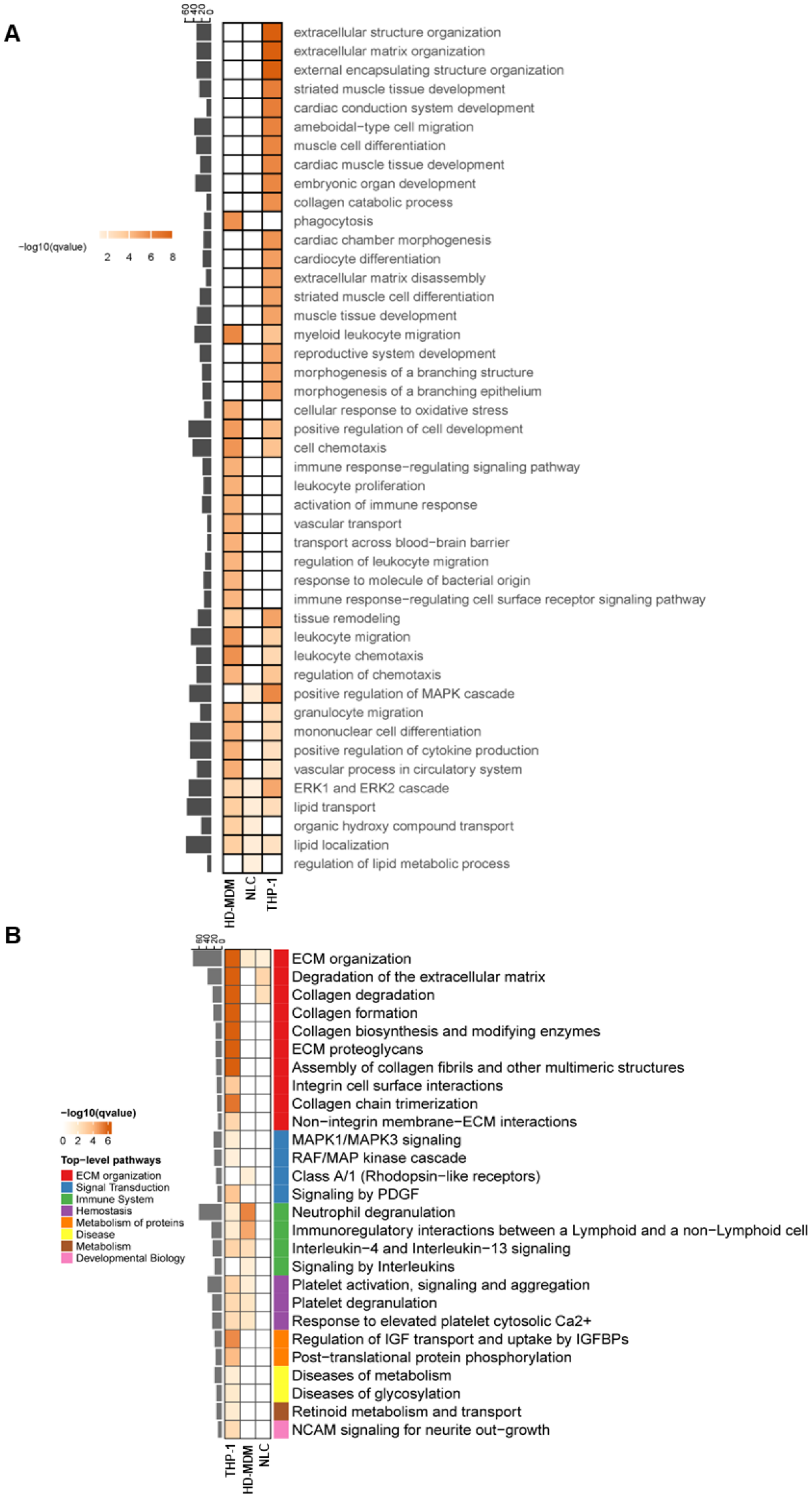
A
-log10(qvalue)
2 4 6 8
60 40 20 0
extracellular structure organization
extracellular matrix organization
external encapsulating structure organization
striated muscle tissue development
cardiac conduction system development
ameboidal−type cell migration
muscle cell differentiation
cardiac muscle tissue development
embryonic organ development
collagen catabolic process
phagocytosis
cardiac chamber morphogenesis
cardiocyte differentiation
extracellular matrix disassembly
striated muscle cell differentiation
muscle tissue development
myeloid leukocyte migration
reproductive system development
morphogenesis of a branching structure
morphogenesis of a branching epithelium
cellular response to oxidative stress
positive regulation of cell development
cell chemotaxis
immune response−regulating signaling pathway
leukocyte proliferation
activation of immune response
vascular transport
transport across blood−brain barrier
regulation of leukocyte migration
response to molecule of bacterial origin
immune response−regulating cell surface receptor signaling pathway
tissue remodeling
leukocyte migration
leukocyte chemotaxis
regulation of chemotaxis
positive regulation of MAPK cascade
granulocyte migration
mononuclear cell differentiation
positive regulation of cytokine production
vascular process in circulatory system
ERK1 and ERK2 cascade
lipid transport
organic hydroxy compound transport
lipid localization
regulation of lipid metabolic process
HD-MDM
NLC
THP-1
B
60 40 20 0
-log10(qvalue)
0 2 4 6
Top−level pathways
ECM organization
Signal Transduction
Immune System
Hemostasis
Metabolism of proteins
Disease
Metabolism
Developmental Biology
ECM organization
Degradation of the extracellular matrix
Collagen degradation
Collagen formation
Collagen biosynthesis and modifying enzymes
ECM proteoglycans
Assembly of collagen fibrils and other multimeric structures
Integrin cell surface interactions
Collagen chain trimerization
Non−integrin membrane−ECM interactions
MAPK1/MAPK3 signaling
RAF/MAP kinase cascade
Class A/1 (Rhodopsin−like receptors)
Signaling by PDGF
Neutrophil degranulation
Immunoregulatory interactions between a Lymphoid and a non−Lymphoid cell
Interleukin−4 and Interleukin−13 signaling
Signaling by Interleukins
Platelet activation, signaling and aggregation
Platelet degranulation
Response to elevated platelet cytosolic Ca2+
Regulation of IGF transport and uptake by IGFBPs
Post−translational protein phosphorylation
Diseases of metabolism
Diseases of glycosylation
Retinoid metabolism and transport
NCAM signaling for neurite out−growth
THP-1
HD-MDM
NLC

(**A**) Heatmap displaying normalized gene expression profiles from RNA-seq data of CLL cells (n = 3), cultured with HD-MDMs, NLCs, and THP-1 macrophages. Columns represent distinct macrophage feeder layer conditions, and rows correspond to genes associated with altered Gene Ontology (GO) Biological Process (BP) terms. Genes were grouped based on enrichment analysis of differentially expressed transcripts. The bar plots adjacent to the heatmap represent the total number of genes contributing to each enriched GO term, -log10(q-value) depicted as color gradient.

(**B**) Heatmap showing normalized gene expression profiles from RNA-seq data of CLL cells (n = 3), cultured with THP-1 macrophages, HD-MDMs, and NLCs. Collums represent individual macrophage feeder layers, and rows correspond to genes associated with significantly enriched Reactome pathways. Genes were grouped based on enrichment analysis of differentially expressed transcripts. The color intensity represents –log10(q-value) for each term. A horizontal bar plot was embedded alongside the heatmap to indicate the gene count associated with each Reactome pathway. Top-level pathways are distinguished by color.

**Figure S10**

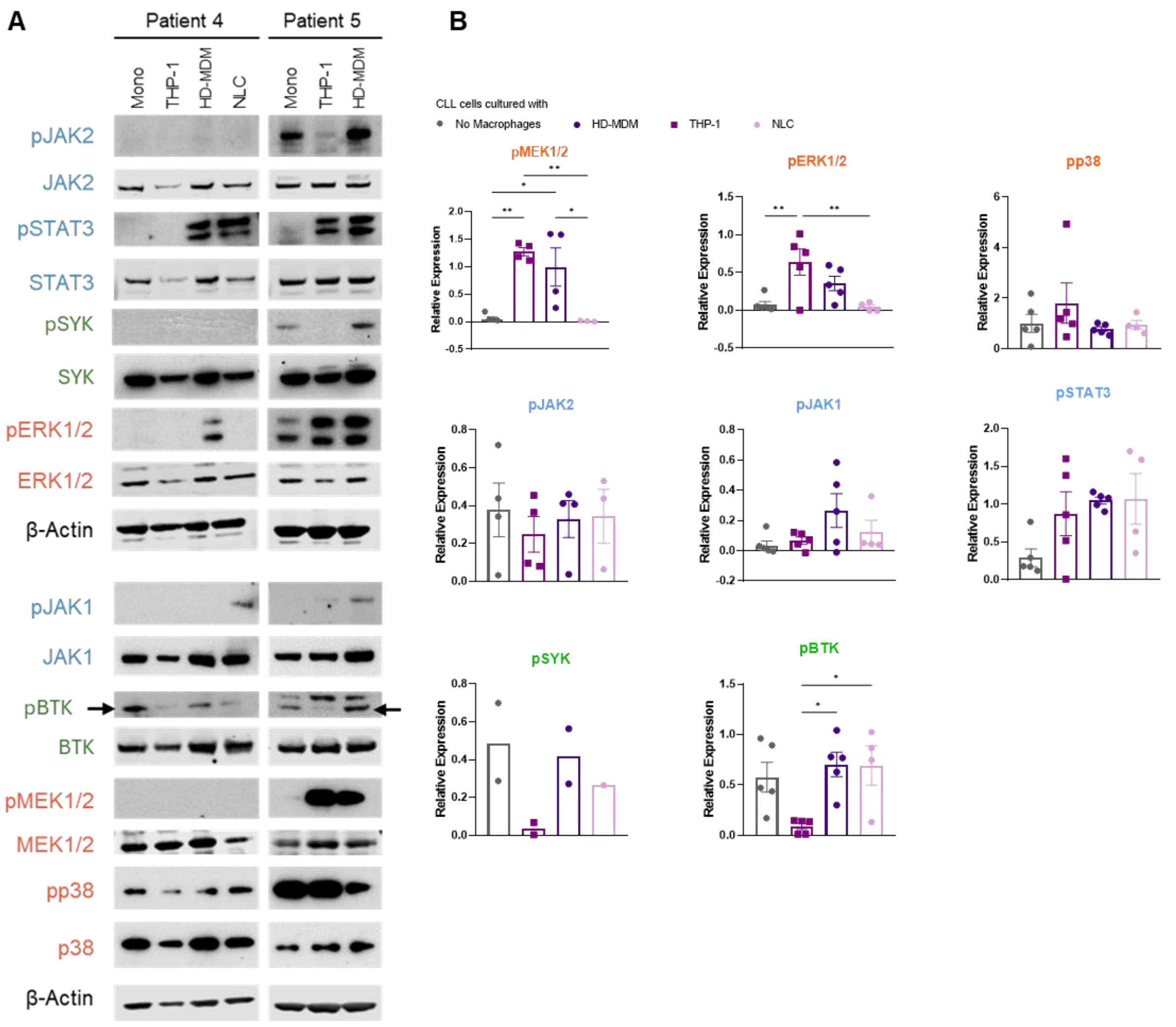

(**A**) CLL cells (n = 2) were lysed after 5 days of monoculture or coculture with THP-1 macrophages, HD-MDMs, or NLCs and analyzed by immunoblotting. Blots were probed for phosphorylated and total forms of JAK2, STAT3, SYK, ERK1/2, JAK1, BTK, MEK1/2, p38MAPK. β-Actin served as a loading control. Members of the JAK/STAT pathway are marked in blue, members of the RAS/MAPK pathway are marked in orange, members of BCR signaling are marked in green.

(**B**) Western Blot quantification of 5 patients (3 in Fig. 2C and 2 in Fig. S10A) by measuring the mean grey value of phospho- compared to total proteins. All proteins were normalized to β-Actin as a loading control. Statistical analysis was performed using an ordinary one-way ANOVA followed by Tukey's multiple comparisons. (p-values are listed in Table S20)

**Figure S11**

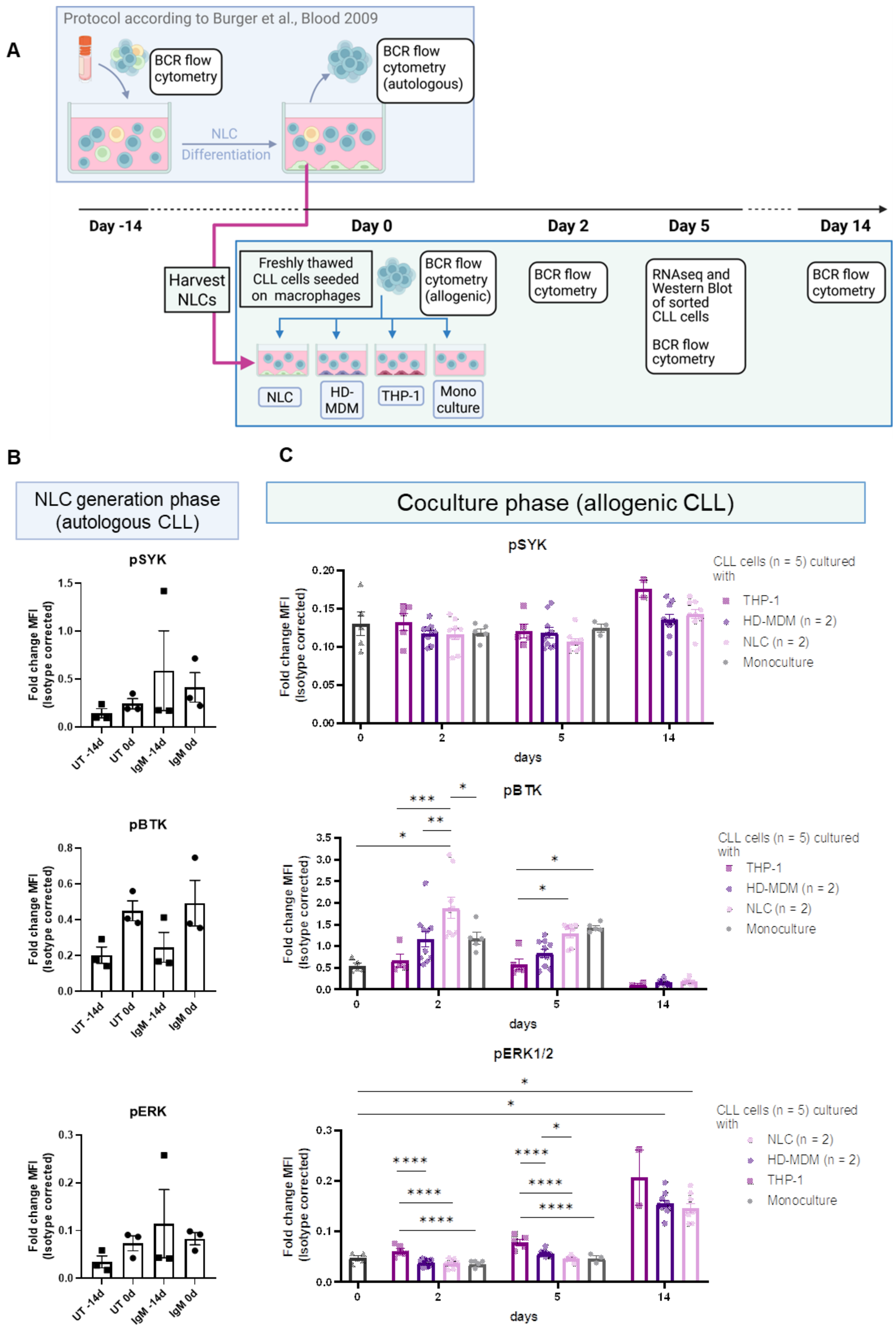
A
Protocol according to Burger et al., Blood 2009
BCR flow cytometry
BCR flow cytometry (autologous)
NLC Differentiation
Day -14
Day 0
Day 2
Day 5
Day 14
Harvest NLCs
Freshly thawed CLL cells seeded on macrophages
BCR flow cytometry (allogenic)
BCR flow cytometry
RNAseq and Western Blot of sorted CLL cells
BCR flow cytometry
BCR flow cytometry
NLC
HD-MDM
THP-1
Mono culture
B
NLC generation phase (autologous CLL)
pSYK
pBTK
pERK
Fold change MFI (Isotype corrected)
UT -14d
UT 0d
IgM -14d
IgM 0d
C
Coculture phase (allogenic CLL)
pSYK
pBTK
pERK1/2
days
CLL cells (n = 5) cultured with
THP-1
HD-MDM (n = 2)
NLC (n = 2)
Monoculture

(**A**) Schematic overview of the experimental workflow for NLC differentiation and CLL analysis used in this study. NLCs were generated by culturing CLL PBMCs for 14 days (from Day -14 to Day 0), following the protocol described by Burger et al. 2009.[10] Then, autologous CLL cells were removed, and newly thawed allogeneic CLL cells were added onto the pre-formed NLCs and other macrophage feeder layers at day 0. The experiments, which were performed at specific days (-14, 0, 2, 5, 14), are indicated in the white boxes.

(**B**) Autologous CLL cells (n = 5) were analyzed by phospho-flow cytometry at Day -14 and Day 0 of NLC differentiation. As controls for BCR activation, CLL cells were stimulated with IgM. Phosphorylation levels of SYK, BTK, and ERK1/2 were assessed and normalized to corresponding total protein levels. No statistical significance was observed with a Kruskal–Wallis test followed by Dunn's multiple comparisons. (p-values are listed in Table S21)

(**C**) Allogeneic CLL cells (n = 5) were analyzed by phospho-flow cytometry at Days 0, 2, 5 and 14 after seeding on macrophage feeder layers. Phosphorylation levels of SYK, BTK, and ERK1/2 were assessed and normalized to corresponding total protein levels. Statistical analysis on different days was performed using a Mixed-effects analysis followed by a Tukey's multiple comparison. (p-values are listed in Table S22) The statistical analysis of different phosphorylation levels compared to the baseline on day 0 was performed using a Kruskal–Wallis test followed by Dunn's multiple comparisons. (p-values are listed in Table S23)

### Supplemental Tables

#### Table S1: Summary of patient IDs used in assays

| Assay | Used Pat. Ids | NLC Ids (if used) |
|---|---|---|
| Viability Coculture | 81 | 81 |
| | 115 | 115 |
| | 182 | 182 |
| | 323 | 323 |
| | 358 | 358 |
| | 430 | 430 |
| | 478 | 478 |
| | 546 | |
| Phagocytosis Assay | 135 | |
| | 430 | |
| | 478 | |
| Phagocytosis Assay (with NLCs, BMDMs and MacCSF1R) | 182 | 182 |
| | 358 | 358 |
| | 457 | 457 |
| ADCP | 182 | |
| Inhibitor Co-Culture | 81 | 317 |
| (Venetoclax/Ibrutinib) | 358 | 478 |
| | 478 | |
| Epcoritamab (CLL PBMCs) | 182 | |
| (THP-1/HD-MDM) | 317 | |
| | 430 | |
| | 457 | |
| | 635 | |
| (NLC) | 331 | 331 |
| | 372 | 372 |
| | 478 | 478 |
| Sequencing coculture | 182 | 545 |
| | 358 | |
| | 420 | |
| Western Blot | 182 | 703 |
| | 504 | 182 |
| | 547 | 182 |
| | 890 | 545 |
| | 904 | |
| BCR phospho-flow | 182 | 182 |
| | 504 | 545 |
| | 547 | |
| | 890 | |
| | 904 | |

**Table S2: Patient information**

| ID | Sex | IGHV-Status |
|---|---|---|
| 81 | male | unmutated |
| 115 | female | mutated |
| 182 | male | mutated |
| 317 | male | mutated |
| 323 | male | unmutated |
| 331 | female | unmutated |
| 358 | female | mutated |
| 372 | male | n.a. |
| 430 | female | unmutated |
| 457 | female | mutated |
| 478 | male | mutated |
| 504 | male | mutated |
| 545 | male | unmutated |
| 546 | female | mutated |
| 547 | n.a. | n.a. |
| 635 | female | unmutated |
| 703 | male | mutated |
| 890 | male | n.a. |
| 904 | male | mutated |

**Table S3: NLC counts**

| Patient NLC-ID | Mean count per well |
|---|---|
| 81 | 1913 |
| 115 | 2599 |
| 182 | 3510 |
| 323 | no NLC could be generated |
| 358 | 1483 |
| 430 | 2281 |
| 478 | 4291 |
| 546 | no NLC could be generated |

**Table S4: List of macrophage-specific genes, excluded from RNAseq data set**

| Gene | Gene synonym | Ensembl | Uniprot |
|---|---|---|---|
| CD300LF | CD300f, CLM1, IGSF13, IREM1, NKIR | ENSG00000186074 | Q8TDQ1 |
| CLEC5A | CLECSF5, MDL-1 | ENSG00000258227 | Q9NY25 |
| CCL18 | AMAC-1, CKb7, DC-CK1, DCCK1, MIP-4, PARC, SCYA18 | ENSG00000275385 | P55774 |
| C1QA | | ENSG00000173372 | P02745 |
| C1QB | | ENSG00000173369 | |
| C1QC | C1QG | ENSG00000159189 | P02747 |
| C3AR1 | AZ3B, C3AR | ENSG00000171860 | Q16581 |
| CD163 | M130, MM130, SCARI1 | ENSG00000177575 | Q86VB7 |
| CD209 | CDSIGN, CLEC4L, DC-SIGN, DC-SIGN1, hDC-SIGN | ENSG00000090659 | Q9NNX6 |
| FCGR1A | CD64, CD64A, FCG1, FcgammaRI, FcgammaRIa, FCGR1 | ENSG00000150337 | P12314 |
| LILRB5 | CD85c, LIR-8, LIR8 | ENSG00000105609 | O75023 |
| MS4A4A | CD20L1, MS4A4, MS4A7 | ENSG00000110079 | Q96JQ5 |
| MS4A4E | | ENSG00000214787 | Q96PG1 |
| MS4A6A | CD20L3, MS4A6 | ENSG00000110077 | Q9H2W1 |
| SIGLEC1 | CD169, dJ1009E24.1, FLJ00051, FLJ00055, FLJ00073, FLJ32150, sialoadhesin, SIGLEC-1, SN | ENSG00000088827 | Q9BZZ2 |
| VSIG4 | CRIg, Z39IG | ENSG00000155659 | Q9Y279 |

**Table S5: List of intersecting DEGs between two or three coculture systems**

| Gene Symbol | expression_type | logFC | adj.P.Val |
|---|---|---|---|
| ANXA1 | all_three | 11,008745 | 0,000103 |
| CD109 | all_three | 11,091306 | 0,000253 |
| CD36 | all_three | 12,734756 | 0,000096 |
| CHI3L1 | all_three | 14,927125 | 0,000096 |
| CST3 | all_three | 10,566287 | 0,000104 |
| CYP1B1 | all_three | 6,460722 | 0,000270 |
| DAB2 | all_three | 12,192983 | 0,000142 |
| DCSTAMP | all_three | 8,390249 | 0,000137 |
| EPB41L3 | all_three | 8,952773 | 0,000274 |
| FNIP2 | all_three | 8,874172 | 0,000142 |
| GAS7 | all_three | 10,285302 | 0,000270 |
| GNA15 | all_three | 9,643599 | 0,000156 |
| GPNMB | all_three | 6,274362 | 0,000487 |
| MGST1 | all_three | 9,558216 | 0,000102 |
| MITF | all_three | 8,042030 | 0,000108 |
| MRAS | all_three | 8,886414 | 0,000102 |
| MYOF | all_three | 11,112679 | 0,000127 |
| NECTIN2 | all_three | 10,076335 | 0,000108 |
| PLAUR | all_three | 9,999834 | 0,000237 |
| PLXDC2 | all_three | 11,168331 | 0,000102 |
| SDC2 | all_three | 13,912632 | 0,000096 |
| SPP1 | all_three | 14,807771 | 0,000096 |
| SUCNR1 | all_three | 10,620669 | 0,000096 |
| TREM2 | all_three | 11,652003 | 0,000096 |
| WDFY3 | all_three | 9,441048 | 0,000129 |
| AARS1 | human_nlc | -1,781436 | 0,019034 |
| ALDH1A1 | human_nlc | 8,532036 | 0,018209 |
| AQP9 | human_nlc | 7,370223 | 0,008897 |
| CYP27A1 | human_nlc | 9,134827 | 0,013305 |
| DLGAP5 | human_nlc | 5,964518 | 0,019034 |
| GAL | human_nlc | 6,592680 | 0,016497 |
| GASK1B | human_nlc | 7,033466 | 0,016593 |
| HS3ST2 | human_nlc | 7,896788 | 0,019263 |
| LINC01484 | human_nlc | -3,211684 | 0,015132 |
| PELATON | human_nlc | 7,424300 | 0,017942 |
| SLC3A2 | human_nlc | -1,529120 | 0,135371 |
| TALDO1 | human_nlc | -1,784968 | 0,018100 |
| TXNRD1 | human_nlc | -1,887484 | 0,013260 |
| ABCC3 | human_thp1 | 9,401698 | 0,000102 |
| ACTN1 | human_thp1 | 8,535062 | 0,001229 |
| ARHGAP10 | human_thp1 | 8,143238 | 0,000293 |
| ATP8B4 | human_thp1 | 10,966864 | 0,000102 |
| C3 | human_thp1 | 7,287801 | 0,000432 |
| CCR1 | human_thp1 | 9,597399 | 0,000242 |
| CD163L1 | human_thp1 | 7,382025 | 0,000129 |
| CDC20 | human_thp1 | 5,692489 | 0,000432 |
| CDK1 | human_thp1 | 7,144604 | 0,000902 |
| CLEC7A | human_thp1 | 8,704923 | 0,000102 |
| CPVL | human_thp1 | 7,704062 | 0,000110 |
| CTSL | human_thp1 | 11,215098 | 0,000141 |
| DAPK1 | human_thp1 | 7,747705 | 0,000525 |
| DHRS3 | human_thp1 | 5,717754 | 0,000382 |
| DYSF | human_thp1 | 8,619653 | 0,000102 |
| EMP1 | human_thp1 | 11,271471 | 0,000706 |
| FCER1G | human_thp1 | 10,326018 | 0,000102 |
| FYB1 | human_thp1 | 10,755592 | 0,000102 |
| GIMAP4 | human_thp1 | 7,048911 | 0,000148 |
| GNLY | human_thp1 | 7,295288 | 0,000499 |
| HNMT | human_thp1 | 9,552891 | 0,000102 |
| ID2 | human_thp1 | 9,658155 | 0,000869 |
| IL1R1 | human_thp1 | 8,100390 | 0,000124 |
| IL7R | human_thp1 | 9,604568 | 0,000251 |
| IRAK3 | human_thp1 | 8,713349 | 0,000102 |
| ITGAL | human_thp1 | 2,742759 | 0,000373 |
| ITGAX | human_thp1 | 6,812090 | 0,000102 |
| KCNE3 | human_thp1 | 8,213640 | 0,000142 |
| KCNMA1 | human_thp1 | 6,710603 | 0,000279 |
| LPAR6 | human_thp1 | 7,367210 | 0,000102 |
| ME1 | human_thp1 | 7,898431 | 0,000268 |
| MMP2 | human_thp1 | 9,610259 | 0,000124 |
| MS4A7 | human_thp1 | 3,384510 | 0,000300 |
| NRP1 | human_thp1 | 11,379010 | 0,000241 |

| | | | |
|---|---|---|---|
| OSBPL1A | human_thp1 | 7,278165 | 0,000826 |
| OTULINL | human_thp1 | 8,095968 | 0,000266 |
| P2RY6 | human_thp1 | 7,278023 | 0,000928 |
| PAG1 | human_thp1 | 8,074120 | 0,000545 |
| PLXNB2 | human_thp1 | 5,868292 | 0,000190 |
| RAB32 | human_thp1 | 7,231532 | 0,000336 |
| RAB7B | human_thp1 | 9,667570 | 0,000148 |
| RNASE1 | human_thp1 | 6,248403 | 0,000783 |
| S100A9 | human_thp1 | 11,130380 | 0,000096 |
| SASH1 | human_thp1 | 9,532203 | 0,001151 |
| SDS | human_thp1 | 6,156989 | 0,000993 |
| SEMA3C | human_thp1 | 6,807409 | 0,000210 |
| SEMA4C | human_thp1 | 5,470746 | 0,000462 |
| SGK1 | human_thp1 | 4,538899 | 0,000601 |
| SH3PXD2B | human_thp1 | 10,570959 | 0,000133 |
| SIGLEC7 | human_thp1 | 6,208716 | 0,000180 |
| SIGLEC9 | human_thp1 | 7,962858 | 0,000164 |
| SLAMF8 | human_thp1 | 9,737599 | 0,000154 |
| SLC8A1 | human_thp1 | 7,121161 | 0,833780 |
| SLCO2B1 | human_thp1 | 9,492572 | 0,833780 |
| SLCO3A1 | human_thp1 | 6,033457 | 0,762689 |
| STAB1 | human_thp1 | 7,714752 | 0,000993 |
| THBD | human_thp1 | 10,417005 | 0,000138 |
| TIMP3 | human_thp1 | 12,481765 | 0,000102 |
| TRPM2 | human_thp1 | 9,066535 | 0,000955 |
| A2M | nlc_thp1 | 8,521344 | 0,018162 |
| AFAP1L1 | nlc_thp1 | 7,399619 | 0,000149 |
| AK1 | nlc_thp1 | 3,646329 | 0,000887 |
| ANO9 | nlc_thp1 | 2,659110 | 0,018944 |
| ATP6V0D2 | nlc_thp1 | 8,665860 | 0,000102 |
| ATP9A | nlc_thp1 | 9,104545 | 0,000102 |
| C1orf21 | nlc_thp1 | 8,578075 | 0,000104 |
| CES1 | nlc_thp1 | 9,459281 | 0,000102 |
| COL6A1 | nlc_thp1 | 12,225211 | 0,000146 |
| COL6A2 | nlc_thp1 | 11,840710 | 0,000148 |
| FMNL1-DT | nlc_thp1 | 3,121146 | 0,000155 |
| GDF15 | nlc_thp1 | 8,844513 | 0,000266 |
| GPC4 | nlc_thp1 | 9,441293 | 0,000339 |
| IQGAP3 | nlc_thp1 | 10,535777 | 0,000102 |
| LINC01010 | nlc_thp1 | 6,019869 | 0,000639 |
| MATK | nlc_thp1 | 6,765756 | 0,010552 |
| MATK | nlc_thp1 | 8,678232 | 0,000102 |
| MMP12 | nlc_thp1 | 10,713862 | 0,000146 |
| MMP19 | nlc_thp1 | 8,403964 | 0,000102 |
| MMP9 | nlc_thp1 | 11,301808 | 0,001189 |
| NCS1 | nlc_thp1 | 9,220182 | 0,000277 |
| RUSC1-AS1 | nlc_thp1 | 2,282422 | 0,000682 |
| SPOCD1 | nlc_thp1 | 9,864315 | 0,000102 |
| TCFL5 | nlc_thp1 | -2,306526 | 0,000142 |
| TM4SF19 | nlc_thp1 | 5,874767 | 0,000246 |

**Table S6: Normalized Enrichment score (NES), p-value, and p-adjusted value for Hallmark GSEA**

| ID | NES | pvalue | p.adjust | qvalue | system |
|---|---|---|---|---|---|
| HALLMARK_ALLOGRAFT_REJECTION | 1,74824874 | 1,7471E-08 | 2,1839E-07 | 1,1034E-07 | Human |
| HALLMARK_ANDROGEN_RESPONSE | 1,46223121 | 0,01640551 | 0,03906074 | 0,01973595 | Human |
| HALLMARK_ANGIOGENESIS | 1,80066077 | 0,00104514 | 0,00522569 | 0,00341045 | NLC |
| HALLMARK_ANGIOGENESIS | 1,70644937 | 3,7761E-05 | 0,00028589 | 0,00013241 | THP1 |
| HALLMARK_APICAL_JUNCTION | 1,5293248 | 0,00166372 | 0,00662747 | 0,00334861 | Human |
| HALLMARK_APICAL_JUNCTION | 1,40515674 | 0,00049916 | 0,00207984 | 0,0009633 | THP1 |
| HALLMARK_APOPTOSIS | 1,52466037 | 0,00208447 | 0,00744453 | 0,00376145 | Human |
| HALLMARK_APOPTOSIS | 1,31171773 | 0,00529947 | 0,01472075 | 0,00681803 | THP1 |
| HALLMARK_CHOLESTEROL_HOMEOSTASIS | 1,52916604 | 0,0004079 | 0,00185408 | 0,00085873 | THP1 |
| HALLMARK_COAGULATION | 1,86759248 | 3,801E-08 | 3,801E-07 | 1,9205E-07 | Human |
| HALLMARK_COAGULATION | 1,79501975 | 1,6211E-05 | 0,00013509 | 8,8164E-05 | NLC |
| HALLMARK_COAGULATION | 1,72586604 | 1,2381E-09 | 1,5476E-08 | 7,1679E-09 | THP1 |
| HALLMARK_COMPLEMENT | 1,76826243 | 4,9764E-09 | 8,2939E-08 | 4,1906E-08 | Human |
| HALLMARK_COMPLEMENT | 1,53508677 | 0,00035089 | 0,00194942 | 0,00127225 | NLC |
| HALLMARK_COMPLEMENT | 1,44069892 | 4,5743E-05 | 0,00028589 | 0,00013241 | THP1 |
| HALLMARK_E2F_TARGETS | 1,31418468 | 0,00127937 | 0,00426457 | 0,00197517 | THP1 |
| HALLMARK_EPITHELIAL_MESENCHYMAL_TRANSITION | 1,7667208 | 9,2995E-07 | 6,6425E-06 | 3,3562E-06 | Human |
| HALLMARK_EPITHELIAL_MESENCHYMAL_TRANSITION | 1,84640397 | 4,4318E-07 | 5,5398E-06 | 3,6154E-06 | NLC |
| HALLMARK_EPITHELIAL_MESENCHYMAL_TRANSITION | 1,72990817 | 7,3112E-15 | 3,6556E-13 | 1,6931E-13 | THP1 |
| HALLMARK_ESTROGEN_RESPONSE_LATE | 1,43673746 | 0,01031244 | 0,02578109 | 0,01302624 | Human |
| HALLMARK_ESTROGEN_RESPONSE_LATE | 1,39967399 | 0,00061823 | 0,00220795 | 0,00102263 | THP1 |
| HALLMARK_G2M_CHECKPOINT | 1,22458442 | 0,02089791 | 0,04543024 | 0,02104137 | THP1 |
| HALLMARK_GLYCOLYSIS | 1,41571269 | 4,5743E-05 | 0,00028589 | 0,00013241 | THP1 |
| HALLMARK_HEDGEHOG_SIGNALING | 1,54221993 | 0,00376911 | 0,01108563 | 0,0051344 | THP1 |
| HALLMARK_HEME_METABOLISM | -1,34622761 | 0,01006137 | 0,0335379 | 0,0218879 | NLC |
| HALLMARK_HYPOXIA | 1,33622038 | 0,00219978 | 0,00687431 | 0,00318389 | THP1 |
| HALLMARK_IL2_STAT5_SIGNALING | 1,40871669 | 0,00705859 | 0,02076055 | 0,01048954 | Human |
| HALLMARK_IL2_STAT5_SIGNALING | 1,30011895 | 0,02881977 | 0,08005492 | 0,05224637 | NLC |
| HALLMARK_IL2_STAT5_SIGNALING | 1,25801339 | 0,01189881 | 0,02952086 | 0,01367282 | THP1 |
| HALLMARK_IL6_JAK_STAT3_SIGNALING | 1,78527812 | 5,2069E-06 | 3,2543E-05 | 1,6443E-05 | Human |
| HALLMARK_IL6_JAK_STAT3_SIGNALING | 1,60180783 | 0,00283873 | 0,01182805 | 0,00771936 | NLC |
| HALLMARK_IL6_JAK_STAT3_SIGNALING | 1,38768385 | 0,00787402 | 0,02072109 | 0,00959714 | THP1 |
| HALLMARK_INFLAMMATORY_RESPONSE | 1,72925759 | 3,252E-07 | 2,71E-06 | 1,3693E-06 | Human |
| HALLMARK_INFLAMMATORY_RESPONSE | 1,48572976 | 0,00233128 | 0,01059674 | 0,00691577 | NLC |
| HALLMARK_INFLAMMATORY_RESPONSE | 1,60976128 | 4,1555E-10 | 6,9258E-09 | 3,2077E-09 | THP1 |
| HALLMARK_INTERFERON_ALPHA_RESPONSE | -1,77887031 | 0,00032704 | 0,00181691 | 0,00091802 | Human |
| HALLMARK_KRAS_SIGNALING_DN | 1,49853465 | 0,00651833 | 0,02507052 | 0,01636181 | NLC |
| HALLMARK_KRAS_SIGNALING_UP | 1,88974911 | 2,6048E-11 | 6,5121E-10 | 3,2903E-10 | Human |
| HALLMARK_KRAS_SIGNALING_UP | 1,70440724 | 1,2249E-05 | 0,00012249 | 7,994E-05 | NLC |
| HALLMARK_KRAS_SIGNALING_UP | 1,64874944 | 6,4829E-11 | 1,6207E-09 | 7,5065E-10 | THP1 |
| HALLMARK_MITOTIC_SPINDLE | 1,38990145 | 0,00813498 | 0,02140784 | 0,01081659 | Human |
| HALLMARK_MITOTIC_SPINDLE | 1,24707788 | 0,01239876 | 0,02952086 | 0,01367282 | THP1 |
| HALLMARK_MTORC1_SIGNALING | -1,44533702 | 0,00308605 | 0,00964391 | 0,00487271 | Human |
| HALLMARK_MTORC1_SIGNALING | -2,34812803 | 1,7963E-13 | 8,9814E-12 | 5,8615E-12 | NLC |
| HALLMARK_MYC_TARGETS_V1 | -1,45150827 | 0,00252489 | 0,00841629 | 0,00425244 | Human |
| HALLMARK_MYC_TARGETS_V1 | -2,18368747 | 8,4907E-12 | 1,4151E-10 | 9,2355E-11 | NLC |
| HALLMARK_MYOGENESIS | 1,54891297 | 0,0015039 | 0,00662747 | 0,00334861 | Human |
| HALLMARK_MYOGENESIS | 1,62953574 | 0,00017115 | 0,0010697 | 0,00069812 | NLC |
| HALLMARK_MYOGENESIS | 1,57793843 | 4,8263E-08 | 4,8263E-07 | 2,2353E-07 | THP1 |
| HALLMARK_NOTCH_SIGNALING | 1,4534263 | 0,0169002 | 0,03840955 | 0,01778969 | THP1 |
| HALLMARK_PANCREAS_BETA_CELLS | 1,62717821 | 0,00811643 | 0,02140784 | 0,01081659 | Human |
| HALLMARK_PROTEIN_SECRETION | -1,84956094 | 0,00011547 | 0,0008248 | 0,00053829 | NLC |
| HALLMARK_REACTIVE_OXYGEN_SPECIES_PATHWAY | -1,47737967 | 0,0229099 | 0,06738206 | 0,04397566 | NLC |
| HALLMARK_SPERMATOGENESIS | 1,49993262 | 0,00058616 | 0,00220795 | 0,00102263 | THP1 |
| HALLMARK_TNFA_SIGNALING_VIA_NFKB | 1,3683057 | 0,00021199 | 0,00105995 | 0,00049092 | THP1 |
| HALLMARK_UNFOLDED_PROTEIN_RESPONSE | -2,48711885 | 6,3758E-12 | 3,1879E-10 | 1,6107E-10 | Human |
| HALLMARK_UNFOLDED_PROTEIN_RESPONSE | -2,72712207 | 7,5307E-13 | 1,8827E-11 | 1,2287E-11 | NLC |
| HALLMARK_UV_RESPONSE_DN | 1,56101281 | 0,00172314 | 0,00662747 | 0,00334861 | Human |
| HALLMARK_UV_RESPONSE_DN | 1,4616588 | 9,6507E-05 | 0,00053615 | 0,00024832 | THP1 |
| HALLMARK_UV_RESPONSE_UP | -1,34182774 | 0,01114446 | 0,03482643 | 0,02272883 | NLC |
| HALLMARK_XENOBIOTIC_METABOLISM | 1,54296668 | 0,00060997 | 0,00304986 | 0,00154098 | Human |
| HALLMARK_XENOBIOTIC_METABOLISM | 1,40125308 | 0,00946711 | 0,0335379 | 0,0218879 | NLC |

**Table S7: Normalized Enrichment score (NES), p-value, and p-adjusted value for GSEA of B cell-specific signatures**

| ID | NES | pvalue | p.adjust | system |
|---|---|---|---|---|
| B cell | 1,24024635 | 0,00331997 | 0,00331997 | THP1 |
| BCR signaling | -1,9121851 | 1,2449E-07 | 1,2449E-07 | Human |
| BCR signaling | -1,67524628 | 7,7815E-06 | 7,7815E-06 | NLC |
| Blimp-1 | 1,56565768 | 0,00590727 | 0,00590727 | Human |
| E2F | 1,33521394 | 0,06537354 | 0,06537354 | Human |
| E2F | 1,22919336 | 0,06230093 | 0,06230093 | THP1 |
| Hypoxia | 1,2597363 | 0,00092999 | 0,00092999 | THP1 |
| IL-10 | 1,36997783 | 0,0353843 | 0,0353843 | Human |
| IL6 | 1,66935286 | 0,00198664 | 0,00198664 | Human |
| IL6 | 1,88977611 | 9,6397E-05 | 9,6397E-05 | NLC |
| IL6 | 1,34316058 | 0,0610752 | 0,0610752 | THP1 |
| JAK | 1,57156817 | 0,01231393 | 0,01231393 | Human |
| JAK | 1,67097008 | 0,00328463 | 0,00328463 | NLC |
| KLF2 | 1,47201218 | 0,00087525 | 0,00087525 | Human |
| Mantle cell lymphoma | 1,60761275 | 0,00132846 | 0,00132846 | THP1 |
| NFkB | 1,31334562 | 0,02108379 | 0,02108379 | Human |
| Notch | 1,25786783 | 0,01235161 | 0,01235161 | Human |
| PAX5 | 1,57309408 | 0,00698047 | 0,00698047 | Human |
| PAX5 | 1,47417109 | 0,00316846 | 0,00316846 | THP1 |
| Proliferation | 1,56924404 | 0,00241758 | 0,00241758 | Human |
| Proliferation | 1,38379547 | 0,03068495 | 0,03068495 | NLC |
| Proliferation | 1,58902321 | 1,8126E-06 | 1,8126E-06 | THP1 |
| Quiescence | 1,2241262 | 0,063967 | 0,063967 | Human |
| RAS | 1,50831465 | 1,3565E-05 | 1,3565E-05 | Human |
| RAS | 1,30735921 | 0,00765077 | 0,00765077 | NLC |
| RAS | 1,47858333 | 2,9733E-10 | 2,9733E-10 | THP1 |
| Serum response | 1,3488754 | 0,00105021 | 0,00105021 | Human |
| SREBP | 1,41699547 | 0,02526618 | 0,02526618 | THP1 |
| STAT3 | 1,47515665 | 0,00571159 | 0,00571159 | Human |
| STAT3 | 1,30135424 | 0,05215054 | 0,05215054 | NLC |
| STAT3 | 1,20390465 | 0,0778693 | 0,0778693 | THP1 |
| T cell calcium signaling | 1,48587677 | 0,01276061 | 0,01276061 | Human |
| T cell cytokine signallng | 1,36696546 | 0,06953757 | 0,06953757 | Human |
| TGF beta | 1,40885943 | 0,05906914 | 0,05906914 | Human |
| TGF beta | 1,5123906 | 0,00063844 | 0,00063844 | THP1 |
| TLR | -1,55074406 | 0,05120948 | 0,05120948 | Human |
| TLR | 1,56470276 | 0,00342754 | 0,00342754 | THP1 |

**Table S8: Primary Western Blot Antibodies**

| Target | Manufacturer | Catalog # |
|---|---|---|
| pJAK2 | Cell Signaling | 3774S |
| JAK2 | Cell Signaling | 3230S |
| pJAK1 | Cell Signaling | 74129T |
| JAK1 | Cell Signaling | 3344T |
| pSTAT3 | Cell Signaling | 9145P |
| STAT3 | Cell Signaling | 9139P |
| pBTK | Cell Signaling | 09120B |
| BTK | Cell Signaling | 85475 |
| pSYK | Cell Signaling | 2710S |
| SYK | Cell Signaling | 2712S |
| pMEK1/2 | Cell Signaling | 9154S |
| MEK1/2 | Cell Signaling | 4694 |
| pp38 | Cell Signaling | 4511S |
| p38 | Cell Signaling | 4511S |
| GAPDH | Cell Signaling | 51745 |
| pERK1/2 | Cell Signaling | 9101S |
| ERK1/2 | Cell Signaling | 9107S |
| β-Actin | Sigma-Aldrich | 065M4837V |

**Table S9: Secondary Western Blot Antibodies**

| Target | Application | Manufacturer | Catalog # |
|---|---|---|---|
| IRDye® 800CW Donkey anti-Rabbit | Li-Cor System | Li-Cor | 926-32213 |
| IRDye® 680LT Donkey anti-Rabbit | Li-Cor System | Li-Cor | 926-68023 |
| IRDye® 800CW Donkey anti-Mouse | Li-Cor System | Li-Cor | 926-32212 |
| IRDye® 680LT Donkey anti-Mouse | Li-Cor System | Li-Cor | 926-68022 |
| Anti-rabbit IgG, HRP-linked Antibody | ECL | Cell Signaling | 7074S |
| Anti-mouse IgG, HRP-linked Antibody | ECL | Cell Signaling | 7076S |

**Table S10: BCR Flow Cytometry Antibodies and Isotypes**

| Target | Clone | Color | Species | Manufacturer | Catalog # |
|---|---|---|---|---|---|
| Annexin V | | FITC | | Immuno Tools | 31490013 |
| Phospho-Syk (Tyr525/526) | C87C1 | PE | human/mouse | Cell Signaling | 6485 |
| Syk (D3Z1E) XP® Rabbit mAb | D3Z1E | Alexa Fluor® 488 | human | Cell Signaling | 13709S |
| ERK1/2 pT202/pY204 Antibody | REA152 | PerCP-Vio® 700 | human/mouse | Miltenyi | 130-103-685 |
| p44/42 MAPK (Erk1/2) (137F5) Rabbit mAb | 137F5 | Alexa Fluor® 647 | diverse | Cell Signaling | 5376S |
| Btk Antibody, anti-human, REAfinity™ | REA367 \| 53/BTK | PE | human | Miltenyi | 130-116-669 |
| BTK [p Tyr223] Antibody - BSA Free | polylconal | Alexa Fluor ® 647 | human/mouse | Novus Biologicals | NBP1-78295 |
| CD19 Antibody, anti-human, REAfinity™ | REA675 \| SJ25-C1 | APC-Vio ® 770 | human | Miltenyi | 130-114-169 |
| REA Control Antibody, human IgG1, REAfinity™ | REA293 \| IS5-21F5 | PerCP-Vio® 700 | human | Miltenyi | 130-113-453 |
| REA Control Antibody (S), human IgG1, REAfinity™ | REA293 \| IS5-21F5 | PE | human | Miltenyi | 130-113-438 |
| REA Control Antibody (S), human IgG1, REAfinity™ | REA293 \| IS5-21F5 | FITC | human | Miltenyi | 130-113-449 |
| REA Control Antibody (S), human IgG1, REAfinity™ | REA293 \| IS5-21F5 | PE-Vio® 770 | human | Miltenyi | 130-113-452 |
| Rabbit (DA1E) mAb IgG XP® Isotype Control | DA1E | Alexa Fluor® 488 | human | Cell Signaling | 2975S |
| Rabbit (DA1E) mAb IgG XP® Isotype Control | DA1E | PE | human | Cell Signaling | 5742S |
| Rabbit (DA1E) mAb IgG XP® Isotype Control | DA1E | Alexa Fluor® 647 | human | Cell Signaling | 2985S |

**Table S11: Reagents**

| Reagent | Manufacturer / Composition |
|---|---|
| ACK lysis buffer | 150 mM NH4Cl, 10 mM KHCO3, 0.1 mM Na2EDTA |
| Antibody diluent | PBS containing 0.04 % Sodium azide, 1 % bovine serum albumin (BSA) |
| Blocking buffer | 1M Tris pH 8.0, 2.5M CaCl2, 5 M NaCl, 0.2 % NP-40, 5 % low-fat milk powder |
| Freezing medium | 50 % RPMI 1640 or DMEM, 40 % FBS, 10 % DMSO |
| GibcoTM RPMI 1640 (Roswell Park Medium Institute) | Thermo Fisher Scientific, Waltham, MA, USA |
| GibcoTM Dulbecco's Modified Eagle Medium (DMEM) | Thermo Fisher Scientific, Waltham, MA, USA |
| GibcoTM DPBS (Dulbecco's phosphate-buffered saline) (14190) | Thermo Fisher Scientific, Waltham, MA, USA |
| GibcoTM Fetal Bovine Serum (FBS) | Thermo Fisher Scientific, Waltham, MA, USA |
| GibcoTM Penicillin-Streptomycin (10,000 U/mL) | Thermo Fisher Scientific, Waltham, MA, USA |
| GlutaMAX™ Supplement | Thermo Fisher Scientific, Waltham, MA, USA |
| M-CSF murine (130-094-129) | Miltenyi Biotec, Bergisch Gladbach, Germany |
| M-CSF human (130-096-491) | Miltenyi Biotec, Bergisch Gladbach, Germany |
| MEM Non-Essential Amino Acids Solution (100X) | Thermo Fisher Scientific, Waltham, MA, USA |
| NuPAGE Antioxidant | Thermo Fisher Scientific, Waltham, MA, USA |
| NuPAGE LDS-Sample buffer (4x) | Thermo Fisher Scientific, Waltham, MA, USA |
| NuPAGE MOPS SDS-Running buffer (20x) | Thermo Fisher Scientific, Waltham, MA, USA |
| NuPAGE Sample Reducing Agent (10x) | Thermo Fisher Scientific, Waltham, MA, USA |
| PBS (phosphate-buffered saline) | 137 mM NaCl, 10 mM Phosphate, 2.7 mM KCl, pH 7.4 |
| Ponceau S staining buffer | 0.1 % Ponceau S, 5 % acetic acid |
| RIPA buffer (10x) | Cell Signaling Technologies, Danvers, MA, USA |
| Sodium Pyruvate (100 mM) | Thermo Fisher Scientific, Waltham, MA, USA |
| TBS-T (Tris-buffered saline with Tween-20) | 50 mM Tris, 150 mM NaCl, 0.1 % Tween 20 |
| Western Blot Transfer Buffer | 25 mM Tris, 192 mM Glycine, 20 % Methanol |
| AnnV-Binding Buffer | BD Biosciences |

**Table S12: Statistics of Fig. 1B (Coculture CLL viability)**

| Šídák's multiple comparisons test | Mean Diff. | 95.00% CI of diff. | Below threshold? | Summary | Adjusted P Value |
|---|---|---|---|---|---|
| Monoculture - HD-MDM (n = 3) | | | | | |
| Day 1 | -16.58 | -28.79 to -4.368 | Yes | ** | 0.0042 |
| Day 3 | -25.56 | -37.77 to -13.35 | Yes | **** | < 0.0001 |
| Day 5 | -42.58 | -54.79 to -30.37 | Yes | **** | < 0.0001 |
| Day 7 | -72.18 | -84.39 to -59.96 | Yes | **** | < 0.0001 |
| Monoculture - NLC | | | | | |
| Day 1 | -8.581 | -17.62 to 0.4586 | No | ns | 0.0691 |
| Day 3 | -10.31 | -19.35 to -1.274 | Yes | * | 0.0195 |
| Day 5 | -21.52 | -30.56 to -12.48 | Yes | **** | < 0.0001 |
| Day 7 | -43.27 | -52.31 to -34.23 | Yes | **** | < 0.0001 |
| Monoculture - THP-1 | | | | | |
| Day 1 | -23.51 | -33.88 to -13.14 | Yes | **** | < 0.0001 |
| Day 3 | -34.98 | -45.35 to -24.61 | Yes | **** | < 0.0001 |
| Day 5 | -47.68 | -58.05 to -37.31 | Yes | **** | < 0.0001 |
| Day 7 | -73.11 | -83.48 to -62.74 | Yes | **** | < 0.0001 |
| Monoculture - BMDM (n = 2) | | | | | |
| Day 1 | -24.46 | -34.05 to -14.87 | Yes | **** | < 0.0001 |
| Day 3 | -38.44 | -48.03 to -28.85 | Yes | **** | < 0.0001 |
| Day 5 | -50.45 | -60.04 to -40.87 | Yes | **** | < 0.0001 |
| Day 7 | -75.87 | -85.45 to -66.28 | Yes | **** | < 0.0001 |
| Monoculture - J774A.1 | | | | | |
| Day 1 | -31.04 | -47.44 to -14.65 | Yes | **** | < 0.0001 |
| Day 3 | -36.48 | -52.88 to -20.08 | Yes | **** | < 0.0001 |
| Day 5 | -45.75 | -62.15 to -29.35 | Yes | **** | < 0.0001 |
| Day 7 | -70.33 | -86.73 to -53.93 | Yes | **** | < 0.0001 |
| Monoculture - MacCsf1r$^{+/+}$ | | | | | |
| Day 1 | -18.55 | -30.34 to -6.763 | Yes | *** | 0.0009 |
| Day 3 | -31.39 | -43.18 to -19.60 | Yes | **** | < 0.0001 |
| Day 5 | -32.35 | -44.14 to -20.56 | Yes | **** | < 0.0001 |
| Day 7 | -55.01 | -66.80 to -43.22 | Yes | **** | < 0.0001 |

**Table S13: Statistics of Fig. 1C (Phagocytosis Assay)**

| Tukey's multiple comparisons test | Mean Diff. | 95.00% CI of diff. | Below threshold? | Summary | Adjusted P Value |
|---|---|---|---|---|---|
| HD-MDM vs. NLC | -8.498 | -54.03 to 37.03 | No | ns | 0.9941 |
| HD-MDM vs. THP-1 | -8.694 | -54.22 to 36.83 | No | ns | 0.9934 |
| HD-MDM vs. BMDM | -23.19 | -68.72 to 22.33 | No | ns | 0.6036 |
| HD-MDM vs. J774A.1 | -56.02 | -101.5 to -10.49 | Yes | * | 0.012 |
| HD-MDM vs. J774A.1 radiated | -16.96 | -62.48 to 28.57 | No | ns | 0.8537 |
| HD-MDM vs. MacCsf1r$^{+/+}$ | -33.23 | -78.75 to 12.30 | No | ns | 0.2333 |
| NLC vs. THP-1 | -0.1963 | -45.72 to 45.33 | No | Ns | > 0.9999 |
| NLC vs. BMDM | -14.7 | -60.22 to 30.83 | No | ns | 0.9173 |
| NLC vs. J774A.1 | -47.52 | -93.05 to -1.991 | Yes | * | 0.0383 |
| NLC vs. J774A.1 radiated | -8.459 | -53.99 to 37.07 | No | ns | 0.9943 |
| NLC vs. MacCsf1r$^{+/+}$ | -24.73 | -70.26 to 20.80 | No | ns | 0.5368 |
| THP-1 vs. BMDM | -14.5 | -60.03 to 31.03 | No | ns | 0.9218 |
| THP-1 vs. J774A.1 | -47.32 | -92.85 to -1.794 | Yes | * | 0.0393 |
| THP-1 vs. J774A.1 radiated | -8.263 | -53.79 to 37.27 | No | ns | 0.9949 |
| THP-1 vs. MacCsf1r$^{+/+}$ | -24.53 | -70.06 to 21.00 | No | ns | 0.5452 |
| BMDM vs. J774A.1 | -32.82 | -78.35 to 12.71 | No | ns | 0.2441 |
| BMDM vs. J774A.1 radiated | 6.237 | -39.29 to 51.77 | No | ns | 0.9989 |
| BMDM vs. MacCsf1r$^{+/+}$ | -10.03 | -55.56 to 35.50 | No | ns | 0.9861 |
| J774A.1 vs. J774A.1 radiated | 39.06 | -6.469 to 84.59 | No | ns | 0.116 |
| J774A.1 vs. MacCsf1r$^{+/+}$ | 22.79 | -22.74 to 68.32 | No | ns | 0.6213 |
| J774A.1 radiated vs. MacCsf1r$^{+/+}$ | -16.27 | -61.80 to 29.26 | No | ns | 0.8751 |

**Table S14: Statistics of Fig. 1D (ADCP)**

| Tukey's multiple comparisons test | Mean Diff | 95,00% CI of diff, | Below threshold? | Summary | Adjusted P Value |
|---|---|---|---|---|---|
| HD-MDM vs. THP-1 | 6,718 | -21,47 to 34,91 | No | ns | 0,7281 |
| HD-MDM vs. BMDM | 1,712 | -10,06 to 13,49 | No | ns | 0,9239 |
| HD-MDM vs. J774A.1 | 18,6 | 12,32 to 24,87 | Yes | ** | 0,0022 |
| HD-MDM vs. MacCsf1r$^{+/+}$ | 9,307 | -1,128 to 19,74 | No | ns | 0,0676 |
| THP-1 vs..BMDM | -5,005 | -40,72 to 30,71 | No | ns | 0,932 |
| THP-1 vs. J774A.1 | 11,88 | -17,17 to 40,94 | No | ns | 0,3766 |
| THP-1 vs. MacCsf1r$^{+/+}$ | 2,59 | -15,50 to 20,68 | No | ns | 0,9274 |
| BMDM vs. J774A.1 | 16,89 | 0,4511 to 33,32 | Yes | * | 0,0465 |
| BMDM vs. MacCsf1r$^{+/+}$ | 7,595 | -10,67 to 25,86 | No | ns | 0,3662 |
| J774A.1 vs. MacCsf1r$^{+/+}$ | -9,292 | -22,61 to 4,024 | No | ns | 0,1251 |

**Table S15: Statistics of Fig. 1E (Inhibitor Assay)**

| Šídák's multiple comparisons test | Mean Diff. | 95.00% CI of diff. | Below threshold? | Summary | Adjusted P Value |
|---|---|---|---|---|---|
| **Ibrutinib 1 µM** | | | | | |
| 24 h | | | | | |
| HD-MDM vs. NLC | 0,0266 | -0,1227 to 0,1759 | No | ns | 0,9624 |
| HD-MDM vs. THP-1 | -0,05904 | -0,2083 to 0,09023 | No | ns | 0,7089 |
| HD-MDM vs. Monoculture | -0,01505 | -0,1643 to 0,1342 | No | ns | 0,9927 |
| NLC vs. THP-1 | -0,08565 | -0,2349 to 0,06362 | No | ns | 0,4182 |
| NLC vs. Monoculture | -0,04165 | -0,1909 to 0,1076 | No | ns | 0,8734 |
| THP-1 vs. Monoculture | 0,044 | -0,1053 to 0,1933 | No | ns | 0,8545 |
| 48 h | | | | | |
| HD-MDM vs. NLC | -0,01797 | -0,1672 to 0,1313 | No | ns | 0,9878 |
| HD-MDM vs. THP-1 | -0,04063 | -0,1899 to 0,1086 | No | ns | 0,8812 |
| HD-MDM vs. Monoculture | 0,02476 | -0,1245 to 0,1740 | No | ns | 0,9692 |
| NLC vs. THP-1 | -0,02266 | -0,1719 to 0,1266 | No | ns | 0,9761 |
| NLC vs. Monoculture | 0,04274 | -0,1065 to 0,1920 | No | ns | 0,8648 |
| THP-1 vs. Monoculture | 0,0654 | -0,08387 to 0,2147 | No | ns | 0,6393 |
| **Venetoclax 5 nM** | | | | | |
| 24 h | | | | | |
| HD-MDM vs. NLC | -0,204 | -0,3640 to -0,04401 | Yes | ** | 0,0068 |
| HD-MDM vs. THP-1 | -0,3545 | -0,5145 to -0,1945 | Yes | **** | <0,0001 |
| HD-MDM vs. Monoculture | -0,2266 | -0,3866 to -0,06654 | Yes | ** | 0,0023 |

| | | | | | |
|---|---|---|---|---|---|
| NLC vs. THP-1 | -0,1505 | -0,3105 to 0,009555 | No | ns | 0,0745 |
| NLC vs. Monoculture | -0,02253 | -0,1825 to 0,1375 | No | ns | 0,9992 |
| THP-1 vs. Monoculture | 0,1279 | -0,03209 to 0,2879 | No | ns | 0,1776 |
| 48 h | | | | | |
| HD-MDM vs. NLC | -0,308 | -0,4680 to -0,1480 | Yes | **** | <0,0001 |
| HD-MDM vs. THP-1 | -0,4673 | -0,6273 to -0,3073 | Yes | **** | <0,0001 |
| HD-MDM vs. Monoculture | -0,3026 | -0,4626 to -0,1426 | Yes | **** | <0,0001 |
| NLC vs. THP-1 | -0,1593 | -0,3193 to 0,0007193 | No | ns | 0,0516 |
| NLC vs. Monoculture | 0,005355 | -0,1547 to 0,1654 | No | ns | >0,9999 |
| THP-1 vs. Monoculture | 0,1647 | 0,004636 to 0,3247 | Yes | * | 0,041 |
| **Ibrutinib 10 µM** | | | | | |
| 24 h | | | | | |
| HD-MDM vs. NLC | 0,09843 | -0,05980 to 0,2567 | No | ns | 0,3961 |
| HD-MDM vs. THP-1 | -0,3777 | -0,5359 to -0,2195 | Yes | **** | <0,0001 |
| HD-MDM vs. Monoculture | -0,02993 | -0,1882 to 0,1283 | No | ns | 0,9944 |
| NLC vs. THP-1 | -0,4761 | -0,6344 to -0,3179 | Yes | **** | <0,0001 |
| NLC vs. Monoculture | -0,1284 | -0,2866 to 0,02987 | No | ns | 0,1521 |
| THP-1 vs. Monoculture | 0,3478 | 0,1896 to 0,5060 | Yes | **** | <0,0001 |
| 48 h | | | | | |
| HD-MDM vs. NLC | 0,02663 | -0,1316 to 0,1849 | No | ns | 0,997 |
| HD-MDM vs. THP-1 | -0,6035 | -0,7617 to -0,4453 | Yes | **** | <0,0001 |

| | | | | | |
|---|---|---|---|---|---|
| HD-MDM vs. Monoculture | 0,03751 | -0,1207 to 0,1957 | No | ns | 0,9819 |
| NLC vs. THP-1 | -0,6301 | -0,7884 to -0,4719 | Yes | **** | <0,0001 |
| NLC vs. Monoculture | 0,01088 | -0,1473 to 0,1691 | No | ns | >0,9999 |
| THP-1 vs. Monoculture | 0,641 | 0,4828 to 0,7992 | Yes | **** | <0,0001 |
| **Venetoclax 10 nM** | | | | | |
| 24 h | | | | | |
| HD-MDM vs. NLC | -0,02222 | -0,2325 to 0,1880 | No | ns | 0,9998 |
| HD-MDM vs. THP-1 | -0,3124 | -0,5226 to -0,1021 | Yes | ** | 0,0024 |
| HD-MDM vs. Monoculture | -0,09193 | -0,3022 to 0,1183 | No | ns | 0,7539 |
| NLC vs. THP-1 | -0,2901 | -0,5004 to -0,07989 | Yes | ** | 0,0046 |
| NLC vs. Monoculture | -0,06971 | -0,2799 to 0,1405 | No | ns | 0,9135 |
| THP-1 vs. Monoculture | 0,2204 | 0,01018 to 0,4307 | Yes | * | 0,0371 |
| 48 h | | | | | |
| HD-MDM vs. NLC | -0,0747 | -0,2849 to 0,1355 | No | ns | 0,8849 |
| HD-MDM vs. THP-1 | -0,3598 | -0,5700 to -0,1495 | Yes | *** | 0,0006 |
| HD-MDM vs. Monoculture | -0,1032 | -0,3134 to 0,1071 | No | ns | 0,6504 |
| NLC vs. THP-1 | -0,2851 | -0,4953 to -0,07483 | Yes | ** | 0,0054 |
| NLC vs. Monoculture | -0,02846 | -0,2387 to 0,1818 | No | ns | 0,9991 |
| THP-1 vs. Monoculture | 0,2566 | 0,04637 to 0,4669 | Yes | * | 0,0126 |

**Table S16: Statistics of Fig. 1F (Bispecific-Ab Killing Assay)**

| Holm-Šídák's multiple comparisons test | Mean Diff. | Below threshold? | Summary | Adjusted P Value |
|---|---|---|---|---|
| Monoculture vs. HD-MDM | 0.2368 | No | ns | 0.1605 |
| Monoculture vs. THP-1 | -0.127 | No | ns | 0.2901 |
| HD-MDM vs. THP-1 | -0.3638 | Yes | * | 0.0186 |

| NLC vs. Monoculture | |
|---|---|
| Paired t test | |
| P value | 0.0633 |
| P value summary | ns |
| Significantly different (P < 0.05)? | No |
| One- or two-tailed P value? | Two-tailed |
| t. df | T = 3.785. df = 2 |
| Number of pairs | 3 |

**Table S17: Statistics of Fig. 2D (BCR Phospho Flow Cytometry)**

| Dunn's multiple comparisons test | Mean rank diff, | Significant? | Summary | Adjusted P Value |
|---|---|---|---|---|
| **pSYK** | | | | |
| NLC (n = 2) vs. HD-MDM (n = 2) | -4,822 | No | ns | >0,9999 |
| NLC (n = 2) vs. THP-1 | -6,622 | No | ns | 0,8082 |
| NLC (n = 2) vs. Monoculture | -10,89 | No | ns | 0,2377 |
| HD-MDM (n = 2) vs. THP-1 | -1,8 | No | ns | >0,9999 |
| HD-MDM (n = 2) vs. Monoculture | -6,067 | No | ns | >0,9999 |
| THP-1 vs. Monoculture | -4,267 | No | ns | >0,9999 |
| **pBTK** | | | | |
| NLC (n = 2) vs. HD-MDM (n = 2) | 9,414 | No | ns | 0,0966 |
| NLC (n = 2) vs. THP-1 | 13,91 | Yes | * | 0,0165 |
| NLC (n = 2) vs. Monoculture | -1,886 | No | ns | >0,9999 |
| HD-MDM (n = 2) vs. THP-1 | 4,5 | No | ns | >0,9999 |
| HD-MDM (n = 2) vs. Monoculture | -11,3 | No | ns | 0,0561 |
| THP-1 vs. Monoculture | -15,8 | Yes | ** | 0,0099 |
| **pERK1/2** | | | | |
| NLC (n = 2) vs. HD-MDM (n = 2) | -9,711 | Yes | * | 0,0465 |
| NLC (n = 2) vs. THP-1 | -17,71 | Yes | *** | 0,0004 |
| NLC (n = 2) vs. Monoculture | -2,111 | No | ns | >0,9999 |
| HD-MDM (n = 2) vs. THP-1 | -8 | No | ns | 0,3945 |
| HD-MDM (n = 2) vs. Monoculture | 7,6 | No | ns | 0,8747 |
| THP-1 vs. Monoculture | 15,6 | Yes | * | 0,0427 |

**Table S18: Statistics of Fig. S5 (M0 M2 Polarization Flow Cytometry)**

| Uncorrected Fisher's LSD | Mean Diff, | 95,00% CI of diff, | Below threshold? | Summary | Individual P Value |
|---|---|---|---|---|---|
| **HD-MDM** | | | | | |
| **M0 MΦ - M2 MΦ** | | | | | |
| CD163 | -4,175 | -19,46 to 11,11 | No | ns | 0,5138 |
| CD204 | -11,16 | -27,94 to 5,624 | No | ns | 0,1481 |
| CD206 | -47,66 | -77,20 to - 18,12 | Yes | ** | 0,0089 |
| **BMDM** | | | | | |
| **M0 MΦ - M2 MΦ** | | | | | |
| CD163 | 95 | -227,8 to 417,8 | No | ns | 0,4561 |
| CD200R | -725,7 | -2113 to 661,5 | No | ns | 0,22 |
| CD206 | -175,3 | -1260 to 909,3 | No | ns | 0,6759 |
| **THP-1** | | | | | |
| **M0 MΦ - M2 MΦ** | | | | | |
| CD36 | -1522 | -9593 to 6549 | No | ns | 0,6752 |
| CD209 | -2841 | -10912 to 5230 | No | ns | 0,4404 |

**Table S19: Statistics of Fig. S6 (XTT Assay)**

| Tukey's multiple comparisons test | Mean Diff, | 95,00% CI of diff, | Below threshold? | Summary | Adjusted P Value |
|---|---|---|---|---|---|
| **NLC low conc.** | | | | | |
| 24 h | | | | | |
| Venetoclax 5 nM vs. ctrl | -0,1715 | -1,072 to 0,7289 | No | ns | 0,8788 |
| Ibrutinib 1 µM vs. ctrl | 0,09103 | -0,8093 to 0,9913 | No | ns | 0,964 |
| 48 h | | | | | |
| Venetoclax 5 nM vs. ctrl | -0,3304 | -1,231 to 0,5700 | No | ns | 0,625 |
| Ibrutinib 1 µM vs. ctrl | 0,1366 | -0,7637 to 1,037 | No | ns | 0,921 |
| **HD-MDM low conc.** | | | | | |
| 24 h | | | | | |
| Venetoclax 5 nM vs. ctrl | 0,0614 | -0,1706 to 0,2934 | No | ns | 0,7989 |
| Ibrutinib 1 µM vs. ctrl | -0,1019 | -0,3339 to 0,1301 | No | ns | 0,5418 |
| 48 h | | | | | |
| Venetoclax 5 nM vs. ctrl | 0,06134 | -0,1707 to 0,2934 | No | ns | 0,7992 |
| Ibrutinib 1 µM vs. ctrl | 0,00537 | -0,2266 to 0,2374 | No | ns | 0,9983 |
| **THP-1 low conc.** | | | | | |
| 24 h | | | | | |
| Venetoclax 5 nM vs. ctrl | -0,03853 | -0,7903 to 0,7132 | No | ns | 0,9898 |
| Ibrutinib 1 µM vs. ctrl | 0,1635 | -0,5882 to 0,9153 | No | ns | 0,833 |
| 48 h | | | | | |
| Venetoclax 5 nM vs. ctrl | 0,1554 | -0,5964 to 0,9072 | No | ns | 0,8477 |
| Ibrutinib 1 µM vs. ctrl | 0,3736 | -0,3781 to 1,125 | No | ns | 0,4084 |

| **HD-MDM high conc.** | | | | | |
|---|---|---|---|---|---|
| 24 h | | | | | |
| Venetoclax 10 nM vs. ctrl | 0,04546 | -0,1684 to 0,2593 | No | ns | 0,8649 |
| Ibrutinib 10 µM vs. ctrl | -0,3903 | -0,6041 to -0,1764 | Yes | *** | 0,0002 |
| 72 h | | | | | |
| Venetoclax 10 nM vs. ctrl | 0,008345 | -0,2055 to 0,2222 | No | ns | 0,9951 |
| Ibrutinib 10 µM vs. ctrl | -0,5153 | -0,7292 to -0,3014 | Yes | **** | <0,0001 |
| **NLC high conc. ONE-WAY ANOVA** | | | | | |
| *Tukey's multiple comparisons test* | *Mean Diff,* | *95,00% CI of diff,* | *Below threshold?* | *Summary* | *Adjusted P Value* |
| Venetoclax 10 nM vs. ctrl | -0,2073 | -0,5686 to 0,1540 | No | ns | 0,3364 |
| Ibrutinib 10 µM vs. ctrl | -0,342 | -0,7033 to 0,01930 | No | ns | 0,0657 |
| **THP-1 high conc. ONE-WAY ANOVA** | | | | | |
| *Tukey's multiple comparisons test* | *Mean Diff,* | *95,00% CI of diff,* | *Below threshold?* | *Summary* | *Adjusted P Value* |
| Venetoclax 10 nM vs. ctrl | -0,01004 | -1,034 to 1,014 | No | ns | 0,9995 |
| Ibrutinib 10 µM vs. ctrl | -0,2642 | -1,288 to 0,7594 | No | ns | 0,7213 |

**Table S20: Statistics of Fig. 2C and S10B (Western Blot Quantification)**

| Tukey's multiple comparisons test | Mean Diff, | 95,00% CI of diff, | Below threshold? | Summary | Adjusted P Value |
|---|---|---|---|---|---|
| **pMEK** | | | | | |
| Monoculture vs. THP-1 | -1,226 | -2,022 to -0,4300 | Yes | ** | 0,0034 |
| Monoculture vs. HD-MDM | -0,9493 | -1,745 to -0,1532 | Yes | * | 0,019 |
| Monoculture vs. NLC | 0,03532 | -0,8245 to 0,8952 | No | ns | 0,9993 |
| THP-1 vs. HD-MDM | 0,2768 | -0,5192 to 1,073 | No | ns | 0,727 |
| THP-1 vs. NLC | 1,261 | 0,4016 to 2,121 | Yes | ** | 0,0049 |
| HD-MDM vs. NLC | 0,9846 | 0,1248 to 1,844 | Yes | * | 0,0241 |
| **pERK** | | | | | |
| Monoculture vs. THP-1 | -0,5702 | -1,004 to -0,1367 | Yes | ** | 0,0086 |
| Monoculture vs. HD-MDM | -0,286 | -0,7196 to 0,1475 | No | ns | 0,2685 |
| Monoculture vs. NLC | 0,02562 | -0,4342 to 0,4854 | No | ns | 0,9985 |
| THP-1 vs. HD-MDM | 0,2841 | -0,1494 to 0,7177 | No | ns | 0,2736 |
| THP-1 vs. NLC | 0,5958 | 0,1360 to 1,056 | Yes | ** | 0,0096 |
| HD-MDM vs. NLC | 0,3117 | -0,1482 to 0,7715 | No | ns | 0,2484 |
| **pp38** | | | | | |
| Monoculture vs. THP-1 | -0,7993 | -2,674 to 1,075 | No | ns | 0,6189 |
| Monoculture vs. HD-MDM | 0,2291 | -1,645 to 2,104 | No | ns | 0,9844 |
| Monoculture vs. NLC | 0,0567 | -1,931 to 2,045 | No | ns | 0,9998 |
| THP-1 vs. HD-MDM | 1,028 | -0,8460 to 2,903 | No | ns | 0,4177 |
| THP-1 vs. NLC | 0,856 | -1,132 to 2,844 | No | ns | 0,6119 |
| HD-MDM vs. NLC | -0,1724 | -2,161 to 1,816 | No | ns | 0,9943 |
| **pJAK2** | | | | | |
| Monoculture vs. THP-1 | 0,1031 | -0,3432 to 0,5495 | No | ns | 0,9083 |
| Monoculture vs. HD-MDM | 0,04113 | -0,4052 to 0,4875 | No | ns | 0,9932 |
| Monoculture vs. NLC | 0,04304 | -0,4304 to 0,5165 | No | ns | 0,9934 |
| THP-1 vs. HD-MDM | -0,06198 | -0,5083 to 0,3844 | No | ns | 0,9775 |
| THP-1 vs. NLC | -0,06007 | -0,5335 to 0,4134 | No | ns | 0,9826 |
| HD-MDM vs. NLC | 0,001909 | -0,4715 to 0,4753 | No | ns | >0,9999 |

| | | | | | |
|---|---|---|---|---|---|
| **pJAK1** | | | | | |
| Monoculture vs. THP-1 | -0,03307 | -0,3140 to 0,2479 | No | ns | 0,986 |
| Monoculture vs. HD-MDM | -0,2322 | -0,5132 to 0,04873 | No | ns | 0,1235 |
| Monoculture vs. NLC | -0,0896 | -0,3876 to 0,2084 | No | ns | 0,8218 |
| THP-1 vs. HD-MDM | -0,1991 | -0,4801 to 0,08180 | No | ns | 0,2164 |
| THP-1 vs. NLC | -0,05653 | -0,3545 to 0,2415 | No | ns | 0,946 |
| HD-MDM vs. NLC | 0,1426 | -0,1554 to 0,4406 | No | ns | 0,5302 |
| **pSTAT3** | | | | | |
| Monoculture vs. THP-1 | -0,5829 | -1,444 to 0,2786 | No | ns | 0,2497 |
| Monoculture vs. HD-MDM | -0,757 | -1,618 to 0,1045 | No | ns | 0,0949 |
| Monoculture vs. NLC | -0,7794 | -1,693 to 0,1343 | No | ns | 0,1082 |
| THP-1 vs. HD-MDM | -0,1741 | -1,036 to 0,6873 | No | ns | 0,9358 |
| THP-1 vs. NLC | -0,1966 | -1,110 to 0,7171 | No | ns | 0,9241 |
| HD-MDM vs. NLC | -0,02245 | -0,9362 to 0,8913 | No | ns | 0,9999 |
| **pSYK** | | | | | |
| Monoculture vs. THP-1 | 0,4517 | -0,5503 to 1,454 | No | ns | 0,3063 |
| Monoculture vs. HD-MDM | 0,06945 | -0,9326 to 1,071 | No | ns | 0,9848 |
| Monoculture vs. NLC | 0,2229 | -1,004 to 1,450 | No | ns | 0,8186 |
| THP-1 vs. HD-MDM | -0,3822 | -1,384 to 0,6198 | No | ns | 0,4033 |
| THP-1 vs. NLC | -0,2288 | -1,456 to 0,9984 | No | ns | 0,8081 |
| HD-MDM vs. NLC | 0,1534 | -1,074 to 1,381 | No | ns | 0,9248 |
| **pBTK** | | | | | |
| Monoculture vs. THP-1 | 0,4869 | -0,03128 to 1,005 | No | ns | 0,0691 |
| Monoculture vs. HD-MDM | -0,1277 | -0,6459 to 0,3905 | No | ns | 0,8914 |
| Monoculture vs. NLC | -0,1151 | -0,6648 to 0,4345 | No | ns | 0,9293 |
| THP-1 vs. HD-MDM | -0,6146 | -1,133 to -0,09644 | Yes | * | 0,0178 |
| THP-1 vs. NLC | -0,602 | -1,152 to -0,05240 | Yes | * | 0,0296 |
| HD-MDM vs. NLC | 0,0126 | -0,5370 to 0,5622 | No | ns | 0,9999 |

#### Table S21: Statistics of Fig. S11B (Phospho-flow NLC generation phase)

| Dunn's multiple comparisons test | Mean rank diff, | Significant? | Summary | Adjusted P Value |
|---|---|---|---|---|
| **pSYK** | | | | |
| UT -14d vs. UT 0d | -3,333 | No | ns | >0,9999 |
| UT -14d vs. IgM -14d | -2,667 | No | ns | >0,9999 |
| UT -14d vs. IgM 0d | -5,333 | No | ns | 0,4202 |
| UT 0d vs. IgM -14d | 0,6667 | No | ns | >0,9999 |
| UT 0d vs. IgM 0d | -2 | No | ns | >0,9999 |
| IgM -14d vs. IgM 0d | -2,667 | No | ns | >0,9999 |
| **pBTK** | | | | |
| UT -14d vs. UT 0d | -6 | No | ns | 0,2492 |
| UT -14d vs. IgM -14d | -1,667 | No | ns | >0,9999 |
| UT -14d vs. IgM 0d | -5 | No | ns | 0,5366 |
| UT 0d vs. IgM -14d | 4,333 | No | ns | 0,8462 |
| UT 0d vs. IgM 0d | 1 | No | ns | >0,9999 |
| IgM -14d vs. IgM 0d | -3,333 | No | ns | >0,9999 |
| **pERK** | | | | |
| UT -14d vs. UT 0d | -3,667 | No | ns | >0,9999 |
| UT -14d vs. IgM -14d | -3,333 | No | ns | >0,9999 |
| UT -14d vs. IgM 0d | -5,667 | No | ns | 0,3255 |
| UT 0d vs. IgM -14d | 0,3333 | No | ns | >0,9999 |
| UT 0d vs. IgM 0d | -2 | No | ns | >0,9999 |
| IgM -14d vs. IgM 0d | -2,333 | No | ns | >0,9999 |

**Table S22: Statistics of Fig. S11C (Phospho-flow Intraday Comparison)**

| Tukey's multiple comparisons test | Predicted (LS) mean diff, | 95,00% CI of diff, | Below threshold? | Summary | Adjusted P Value |
|---|---|---|---|---|---|
| **pSYK** | | | | | |
| Day 2 | | | | | |
| NLC (n = 2) vs. HD-MDM (n = 2) | -0,001194 | -0,02249 to 0,02010 | No | ns | 0,9988 |
| NLC (n = 2) vs. THP-1 | -0,01509 | -0,04082 to 0,01065 | No | ns | 0,4105 |
| NLC (n = 2) vs. Monoculture | -0,002317 | -0,02807 to 0,02344 | No | ns | 0,9951 |
| HD-MDM (n = 2) vs. THP-1 | -0,0139 | -0,03916 to 0,01137 | No | ns | 0,467 |
| HD-MDM (n = 2) vs. Monoculture | -0,001123 | -0,02641 to 0,02416 | No | ns | 0,9994 |
| THP-1 vs. Monoculture | 0,01277 | -0,01635 to 0,04189 | No | ns | 0,6502 |
| Day 5 | | | | | |
| NLC (n = 2) vs. HD-MDM (n = 2) | -0,01174 | -0,03304 to 0,009555 | No | ns | 0,4648 |
| NLC (n = 2) vs. THP-1 | -0,01559 | -0,04132 to 0,01015 | No | ns | 0,3818 |
| NLC (n = 2) vs. Monoculture | -0,02012 | -0,05060 to 0,01036 | No | ns | 0,3066 |
| HD-MDM (n = 2) vs. THP-1 | -0,003843 | -0,02911 to 0,02142 | No | ns | 0,9773 |
| HD-MDM (n = 2) vs. Monoculture | -0,008376 | -0,03846 to 0,02171 | No | ns | 0,88 |
| THP-1 vs. Monoculture | -0,004532 | -0,03791 to 0,02884 | No | ns | 0,9836 |
| Day 14 | | | | | |
| NLC (n = 2) vs. HD-MDM (n = 2) | 0,008001 | -0,01588 to 0,03188 | No | ns | 0,6719 |
| NLC (n = 2) vs. THP-1 | -0,03287 | -0,1318 to 0,06610 | No | ns | 0,2586 |
| HD-MDM (n = 2) vs. THP-1 | -0,04087 | -0,1279 to 0,04616 | No | ns | 0,1791 |

| | | | | | |
|---|---|---|---|---|---|
| **pBTK** | | | | | |
| Day 2 | | | | | |
| NLC (n = 2) vs. HD-MDM (n = 2) | 0,7152 | 0,1610 to 1,269 | Yes | ** | 0,0066 |
| NLC (n = 2) vs. THP-1 | 1,232 | 0,5194 to 1,945 | Yes | *** | 0,0002 |
| NLC (n = 2) vs. Monoculture | 0,6987 | 0,02591 to 1,371 | Yes | * | 0,0391 |
| HD-MDM (n = 2) vs. THP-1 | 0,5168 | -0,1844 to 1,218 | No | ns | 0,2165 |
| HD-MDM (n = 2) vs. Monoculture | -0,01654 | -0,6772 to 0,6441 | No | ns | 0,9999 |
| THP-1 vs. Monoculture | -0,5333 | -1,332 to 0,2649 | No | ns | 0,2958 |
| Day 5 | | | | | |
| NLC (n = 2) vs. HD-MDM (n = 2) | 0,492 | -0,09292 to 1,077 | No | ns | 0,1273 |
| NLC (n = 2) vs. THP-1 | 0,752 | 0,05365 to 1,450 | Yes | * | 0,0303 |
| NLC (n = 2) vs. Monoculture | -0,08681 | -0,7851 to 0,6115 | No | ns | 0,9873 |
| HD-MDM (n = 2) vs. THP-1 | 0,2599 | -0,4007 to 0,9206 | No | ns | 0,7223 |
| HD-MDM (n = 2) vs. Monoculture | -0,5788 | -1,240 to 0,08183 | No | ns | 0,1049 |
| THP-1 vs. Monoculture | -0,8388 | -1,602 to -0,07591 | Yes | * | 0,026 |
| Day 14 | | | | | |
| NLC (n = 2) vs. HD-MDM (n = 2) | 0,02588 | -0,04985 to 0,1016 | No | ns | 0,6516 |
| NLC (n = 2) vs. THP-1 | 0,0659 | -0,1064 to 0,2382 | No | ns | 0,3302 |
| HD-MDM (n = 2) vs. THP-1 | 0,04001 | -0,1760 to 0,2560 | No | ns | 0,5673 |
| **pERK** | | | | | |
| Day 2 | | | | | |

| | | | | | |
|---|---|---|---|---|---|
| NLC (n = 2) vs. HD-MDM (n = 2) | -0,002754 | -0,01271 to 0,007202 | No | ns | 0,882 |
| NLC (n = 2) vs. THP-1 | -0,02488 | -0,03696 to -0,01279 | Yes | **** | <0,0001 |
| NLC (n = 2) vs. Monoculture | 0,0003548 | -0,01173 to 0,01244 | No | ns | 0,9998 |
| HD-MDM (n = 2) vs. THP-1 | -0,02212 | -0,03399 to -0,01025 | Yes | **** | <0,0001 |
| HD-MDM (n = 2) vs. Monoculture | 0,003109 | -0,008760 to 0,01498 | No | ns | 0,8977 |
| THP-1 vs. Monoculture | 0,02523 | 0,01153 to 0,03894 | Yes | **** | <0,0001 |
| Day 5 | | | | | |
| NLC (n = 2) vs. HD-MDM (n = 2) | -0,01027 | -0,02022 to -0,0003126 | Yes | * | 0,0409 |
| NLC (n = 2) vs. THP-1 | -0,0323 | -0,04438 to -0,02021 | Yes | **** | <0,0001 |
| NLC (n = 2) vs. Monoculture | 0,0008829 | -0,01294 to 0,01470 | No | ns | 0,9982 |
| HD-MDM (n = 2) vs. THP-1 | -0,02203 | -0,03390 to -0,01016 | Yes | **** | <0,0001 |
| HD-MDM (n = 2) vs. Monoculture | 0,01115 | -0,002478 to 0,02478 | No | ns | 0,1442 |
| THP-1 vs. Monoculture | 0,03318 | 0,01792 to 0,04843 | Yes | **** | <0,0001 |
| Day 14 | | | | | |
| NLC (n = 2) vs. HD-MDM (n = 2) | -0,009079 | -0,03746 to 0,01931 | No | ns | 0,6921 |
| NLC (n = 2) vs. THP-1 | -0,06061 | -0,9744 to 0,8532 | No | ns | 0,6549 |
| HD-MDM (n = 2) vs. THP-1 | -0,05153 | -0,9970 to 0,8939 | No | ns | 0,7142 |

**Table S23: Statistics of Fig. S11C (BCR Baseline Comparison)**

| Dunn's multiple comparisons test | Mean rank diff, | Significant? | Summary | Adjusted P Value |
|---|---|---|---|---|
| **pSYK** | | | | |
| Baseline Day 0 vs. NLC Day 2 | 9,467 | No | ns | >0,9999 |
| Baseline Day 0 vs. HD-MDM Day 2 | 9,2 | No | ns | >0,9999 |
| Baseline Day 0 vs. THP-1 Day 2 | -5,6 | No | ns | >0,9999 |
| Baseline Day 0 vs. Monoculture Day 2 | 7 | No | ns | >0,9999 |
| Baseline Day 0 vs. NLC Day 5 | 23,13 | No | ns | 0,8975 |
| Baseline Day 0 vs. HD-MDM Day 5 | 10,1 | No | ns | >0,9999 |
| Baseline Day 0 vs. THP-1 Day 5 | 7,2 | No | ns | >0,9999 |
| Baseline Day 0 vs. Monoculture Day 5 | -0,8667 | No | ns | >0,9999 |
| Baseline Day 0 vs. NLC Day 14 | -18,09 | No | ns | >0,9999 |
| Baseline Day 0 vs. HD-MDM Day 14 | -10,3 | No | ns | >0,9999 |
| Baseline Day 0 vs. THP-1 Day 14 | -36,2 | No | ns | 0,7617 |
| **pBTK** | | | | |
| Baseline Day 0 vs. NLC Day 2 | -37,33 | Yes | * | 0,0437 |
| Baseline Day 0 vs. HD-MDM Day 2 | -21,8 | No | ns | 0,9543 |
| Baseline Day 0 vs. THP-1 Day 2 | -4 | No | ns | >0,9999 |
| Baseline Day 0 vs. Monoculture Day 2 | -23,2 | No | ns | >0,9999 |
| Baseline Day 0 vs. NLC Day 5 | -30,43 | No | ns | 0,2787 |
| Baseline Day 0 vs. HD-MDM Day 5 | -10,1 | No | ns | >0,9999 |
| Baseline Day 0 vs. THP-1 Day 5 | 0,6 | No | ns | >0,9999 |

| | | | | |
|---|---|---|---|---|
| Baseline Day 0 vs. Monoculture Day 5 | -34,6 | No | ns | 0,2042 |
| Baseline Day 0 vs. NLC Day 14 | 18,38 | No | ns | >0,9999 |
| Baseline Day 0 vs. HD-MDM Day 14 | 21,1 | No | ns | >0,9999 |
| Baseline Day 0 vs. THP-1 Day 14 | 26 | No | ns | >0,9999 |
| **pERK** | | | | |
| Baseline Day 0 vs. NLC Day 2 | 17,24 | No | ns | >0,9999 |
| Baseline Day 0 vs. HD-MDM Day 2 | 13,8 | No | ns | >0,9999 |
| Baseline Day 0 vs. THP-1 Day 2 | -17 | No | ns | >0,9999 |
| Baseline Day 0 vs. Monoculture Day 2 | 19,6 | No | ns | >0,9999 |
| Baseline Day 0 vs. NLC Day 5 | 1,689 | No | ns | >0,9999 |
| Baseline Day 0 vs. HD-MDM Day 5 | -14,5 | No | ns | >0,9999 |
| Baseline Day 0 vs. THP-1 Day 5 | -26 | No | ns | 0,9275 |
| Baseline Day 0 vs. Monoculture Day 5 | 0,8 | No | ns | >0,9999 |
| Baseline Day 0 vs. NLC Day 14 | -37,87 | Yes | * | 0,048 |
| Baseline Day 0 vs. HD-MDM Day 14 | -41,2 | Yes | * | 0,0174 |
| Baseline Day 0 vs. THP-1 Day 14 | -45,7 | No | ns | 0,24 |
